# Plant-on-Chip: core morphogenesis processes in the tiny plant *Wolffia australiana*

**DOI:** 10.1101/2022.04.16.488569

**Authors:** Feng Li, Jing-Jing Yang, Zong-Yi Sun, Lei Wang, Le-Yao Qi, A Sina, Yi-Qun Liu, Hong-Mei Zhang, Lei-Fan Dang, Shu-Jing Wang, Chun-Xiong Luo, Wei-Feng Nian, Seth O’Conner, Long-Zhen Ju, Wei-Peng Quan, Xiao-Kang Li, Chao Wang, De-Peng Wang, Han-Li You, Zhu-Kuan Cheng, Jia Yan, Fu-Chou Tang, De-Chang Yang, Chu-Wei Xia, Ge Gao, Yan Wang, Bao-Cai Zhang, Yi-Hua Zhou, Xing Guo, Sun-Huan Xiang, Huan Liu, Tian-Bo Peng, Xiao-Dong Su, Yong Chen, Qi Ouyang, Dong-Hui Wang, Da-Ming Zhang, Zhi-Hong Xu, Hong-Wei Hou, Shu-Nong Bai, Ling Li

## Abstract

A plant can be thought of as a colony comprising numerous growth buds, each developing to its own rhythm. Such lack of synchrony impedes efforts to describe core principles of plant morphogenesis, dissect the underlying mechanisms, and identify regulators. Here, we use the tiniest known angiosperm to overcome this challenge and provide an ideal model system for plant morphogenesis. We present a detailed morphological description of the monocot *Wolffia australiana*, as well as high-quality genome information. Further, we developed the Plant-on-Chip culture system and demonstrate the application of advanced technologies such as snRNA-seq, protein structure prediction, and gene editing. We provide proof-of-concept examples that illustrate how *W. australiana* can open a new horizon for deciphering the core regulatory mechanisms of plant morphogenesis.

**Significance:** What is the core morphogenetic process in angiosperms, a plant like a tree indeterminately growing, or a bud sequentially generating limited types of organs? *Wolffia australiana*, one of the smallest angiosperms in the world may help to make a distinction. Wolffia plantlet constitutes of only three organs that are indispensable to complete life cycle: one leaf, one stamen and one gynoecium. Before the growth tip is induced to flower, it keeps branching from the leaf axil and the branches separate from the main plantlet. Here we present a high-quality genome of *W. australiana*, detailed morphological description, a Plant-on-Chip cultural system, and some principle-proof experiments, demonstrating that *W. australiana* is a promising model system for deciphering core developmental program in angiosperms.

## Introduction

What are the core morphogenetic processes required for a multicellular organism to complete its life cycle? For most species in the animal kingdom, embryogenesis plays such a core role, with all organs initiated at that stage. By contrast, in most species in the plant kingdom, organs and tissues are produced sequentially. Plant development starts, like that of animals, with the formation of a zygote, whose number and types of organs are limited. However, plants then go on to produce an indeterminate number of organs such as leaves, roots and stems before they produce spores and initiate the gametophyte for formation of haploid gametes. Also in contrast to animals, during the process of completing the life cycle, in addition to the growth tip derived from the zygote, many plants can produce new growth tips in the axillary buds.

C. H. Waddington (1) pointed out that each apical meristem “gives rise to a whole new cycle of growth and development.” Since a plant comprises many branches derived from lateral growth, each plant may be seen as a colony of buds (or meristems) supporting growth, in essence rather comparable to the outer shell of a coral colony than to an individual worm, bird, or cat. Notably, unlike the broad synchronization of polyps inhabiting coral structures in their developmental process on an annual rhythm, plant branches or buds undergo their development independently of one another. For example, in the perennial plant alpine rock cress (*Arabis alpina*), a close relative of the model plant Arabidopsis (*Arabidopsis thaliana*), a subset of meristems will transition to their reproductive stage under conditions conducive to flowering, while other meristems will remain vegetative (2). Likewise, apple (*Malus domestica*) trees carry vegetative buds and floral buds in various developmental stages simultaneously to support vegetative growth and fruit harvest each year.

While the morphogenetic strategy combining apical growth and branching rendered advantages for plants as sessile and photoautotrophic eukaryotes, this morphogenetic strategy can make it difficult to elucidate the core morphogenetic processes due to a general lack of synchronization between meristems. We reasoned that truly systematic dissection of the mechanisms driving the core principles of plant morphogenesis requires a plant species with simple branching, few organs, and clearly distinguishable morphological boundaries for empirical investigations. We propose here that the genus *Wolffia* provides such an ideal plant (Fig. 1).

**Fig. 1.**
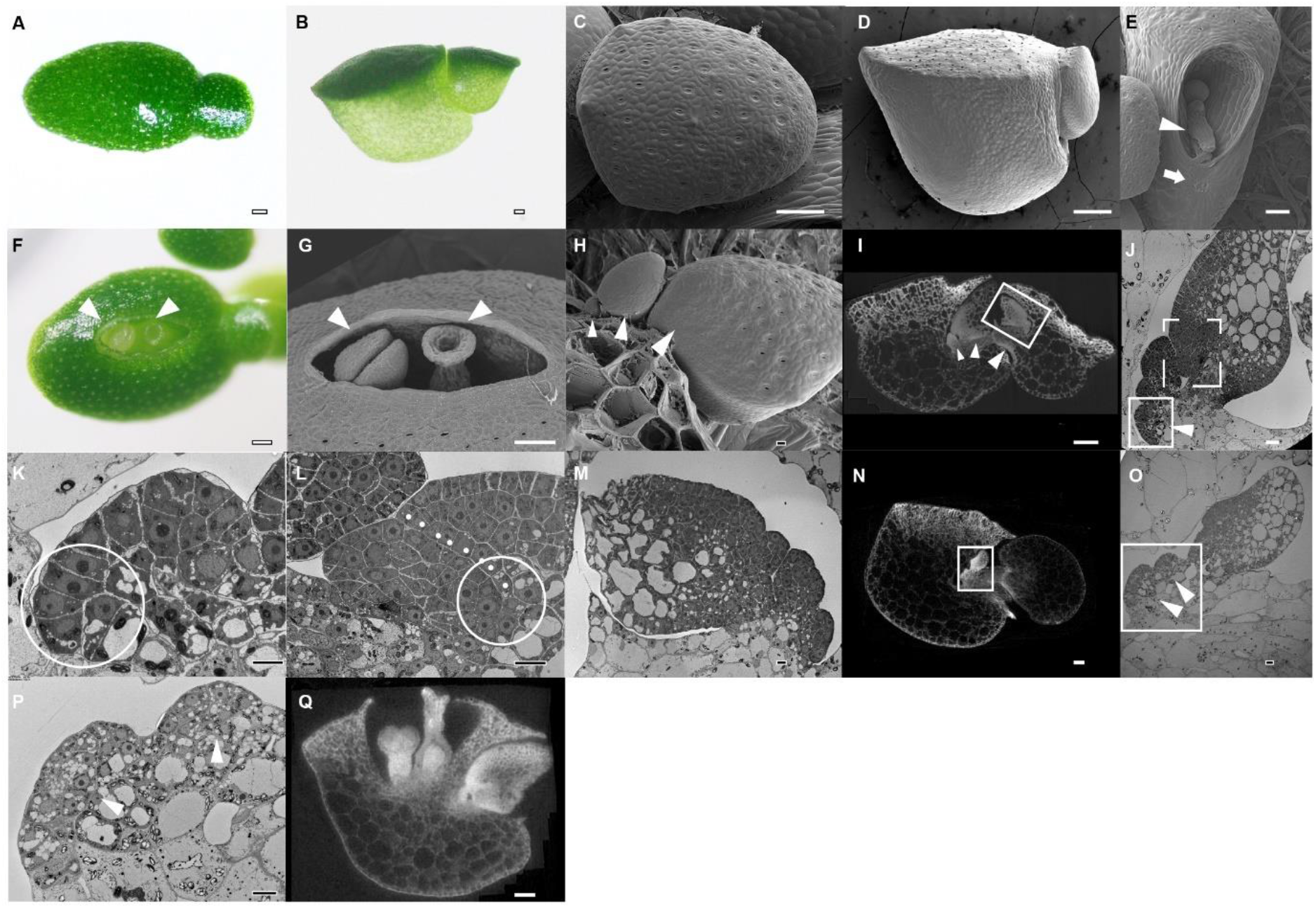
Morphological description of *W. australiana*. (*A*) Top view of a *W. australiana* plantlet, as seen under a dissecting microscope; a branch is protruding to the right. (*B*) Side view of a *W. australiana* plantlet, showing a boat-shaped leaf with dark green cells at the “deck” and light green cells at the “hull.” (*C*) Top view of a *W. australiana* plantlet by cryo-SEM; note the presence of stomata. (*D*) Side view of a *W. australiana* plantlet by cryo-SEM; no stomata were observed. (*E*) View from the hole from which branches abscise out, showing the remaining petiole (arrowhead). A scar (arrow) forms on the boat-shaped leaf and indicates prior branch abscission. (*F*) Top view of a *W. australiana* plantlet under a dissecting microscope, showing the stigma and stamen (arrowheads) protruding from the crack on the “deck.” (*G*) Top view of the crack region of a *W. australiana* plantlet, as seen by cryo-SEM, showing a stigma (right, arrowhead) and a stamen (left, arrowhead). (*H*) After peeling the deck, three young leaves (the biggest one has developed into a branch) are aligned sequentially, as indicated by three arrowheads. (*I*) Computed tomography (CT) image showing the alignment of leaves (arrowheads) and how the biggest leaf has developed into a branch before abscission (the rectangle shows new leaves produced from the biggest leaf). (*J*) Transmission electron microscopy (TEM) section of the leaf primordia and the region including the growth tip (rectangle with arrowhead). (*K*) Zoomed-in region indicated by the solid rectangle shown in *J*. The circle highlights the cells with big nuclei and dense cytoplasm, possible including the growth tip cell(s). (*L)* Zoomed-in region indicated by the dashed rectangle in *J* with adjusted orientation. The dotted line indicates the border of the fast-growing region of the primordium leaf that overlaps with the slow-growing region. The circle indicates the junction where a growth tip of the primordium leaf might initiate *de novo*, which allows a primordium leaf to become a new branch. (*M*) TEM section of the growth region of the branch (prepared with the CT sample, corresponding to the corresponding rectangle in Fig. 1I). (*N*) CT image showing a region of the growth tip of a plantlet under flower induction conditions. The rectangle highlights the growth region for further observation. (*O*) TEM section of the growth region (prepared with the CT sample, corresponding to the corresponding rectangle in Fig. 1***N***). Two bumps (arrowheads) arise from the innermost region of the cavity. (*P*) Further enlargement of the region highlighted in Fig. 1*O*. Arrowheads indicate cells in the bumps that are morphologically different from those shown in Fig. 1*K*. (*Q*) CT image showing a gynoecium (right) and a stamen (left) inside the plantlet, possibly derived from the two bumps observed in Fig. 1*P*. Bars = 100 µm (*A, B, C, D, E, F, G, I, N, Q*) and 10 µm (*H, J, K, L, M, O, P*).

*Wolffia* is a genus of aquatic plants. It is the simplest and smallest known angiosperm in the world (3, 4). Each *Wolffia australiana* plantlet consists of a single boat-like leaf with a diameter of 1 mm, with a few axillary buds wrapped around the base of the leaf, but no root. After the axillary buds grow, they separate (abscise) from the main plantlet. Under stress conditions, the plantlet produces one stamen and one gynoecium, which arise vertically upward from a hole generated in the center of the “deck” of the boat-like leaf. The stamen consists of two locules containing numerous pollen grains, while the gynoecium contains a single ovule (5, 6); Fig. 1). Therefore, the *Wolffia* plantlet harbors a minimal set of core organs needed for an angiosperm to complete its life cycle, and not much more.

What insights might be gleaned using the *Wolffia* plantlet as a model system? Work on the tiny angiosperm over the past 60 years provides some clues. For instance, the transition from vegetative branching to floral organs only takes a few days, thus offering a unique opportunity to decipher the mechanisms of cell fate change along a precise spatiotemporal pattern (7, 8). *Wolffia* as a model system may thus unlock the developmental programs underlying plant morphogenesis.

The shoot apical meristem (SAM) of *Wolfia* plantlets is also much simpler than that of other spermatophytes. While the shoot tip is made up of a cell cluster in gymnosperms and angiosperms, a single cell or a few cells are sufficient to carry out the functions of a growth tip, as in the haploid moss *Physcomitrium* (*Physcomitrella*) *patens*, or in the diploid pteridophytes such as *Selaginella* or *Adiantum*. Considering that the entire *Wolffia* plantlet is only 1 mm in diameter, and based on observations that revealed too few cells at the tip to organize into a higher-order structures as that seen in angiosperms (6, 9-13), the *Wolffia* growth tip holds the promise of a much simplified organization with no tunica-corpus structure that can produce leaves, axillary buds, and finally the stamen and gynoecium.

All species in the *Wolffia* genus lack a clear root, while other closely related duckweed species, such as *Lemna minor, Lemna gibba*, and *Spirodela polyrhiza*, all have roots. The absence of the root is therefore unlikely to represent an adaptation to an aquatic environment. Using the rich information available on root development in land plants such as Arabidopsis, rice (*Oryza sativa*), and maize (*Zea mays*), it should be possible to generate testable hypotheses to mechanistically explain the absence of roots in *Wolffia* and contribute to our understanding of the origin(s) and morphogenesis of roots in angiosperms.

Perhaps the greatest advantage of *Wolffia* plantlets is that they only carry a single leaf, as each new branch will bud off as a separate plantlet. This growth pattern thus provides unique opportunities to continuously observe the same plantlet for its entire life cycle under a microscope.

The simplicity of the *Wolffia* genus and its duckweeds relatives has attracted much interest and has led to efforts to develop this tiny plant(let) into an experimental system (3-5, 14, 15). However, key tools are currently lacking to propel the *Wolffia* genus as a model system to investigate core principles of the plant life cycle. Here, we report our efforts in developing *W. australiana* as a model system. We sequenced the genome of this species, which will complement other genome sequences from this species recently published (16, 17) and set up a “Plant-on-Chip (PoC)” culture system with which to observe morphological characteristics, particularly in the diploid phase. We also exploited the genomic information we generated to analyze the possible mechanisms behind unique morphological traits seen in *W. australiana* such as the lack of a root or a vasculature and the fast transition from vegetative to reproductive growth. Furthermore, we demonstrated the feasibility of obtaining transgenic plantlets, single-nucleus RNA-sequencing (snRNA-seq) and protein structure prediction. We believe that this new PoC system will serve as a platform for the dissection of core principles underlying plant morphogenesis and predict that our PoC system will open new avenues in plant biology research.

## Results

### Morphological Uniqueness: Growth Tip, Branch, and Floral Organs

A typical plantlet (previously termed a “frond”) of *W. australiana* has one boat-shaped leaf with a branch (previously called a daughter frond) on the side (6) (Fig. 1 *A* and *B*). Our detailed observations (below) revealed that frond is not the correct term for these structures. We therefore refer to them as plantlets and branches instead of fronds and daughter fronds, respectively, hereafter.

Close observations revealed that the deck part of the boat-shaped leaf is relatively dark green with smaller cells, while the hull part was relatively light green with larger cells (Fig. 1*B*). Using cryo-scanning electron microscopy, we observed stomata on the surface of the deck but not on the hull surface (Fig. 1 *C* and *D*). We also noticed the presence of a scar near the hole from where the branched plantlets continuously protrude out (Fig. 1*E*), corresponding to plantlet abscission from the petiole linking the branching plantlet. The hole and scar provided clear markers to orientate a plantlet. Under inductive conditions (see below), we observed the emergence of a crack in the center of the deck, perpendicularly to the hole from which branching plantlets protrude. The floral organs, one stamen and one gynoecium, rose from the crack (Fig. 1 *F* and *G*; video at www.wolffiapond.net, Username and Password: waus).

After peeling off the deck, we noticed three consecutive plantlets (on growing branches) of decreasing size (Fig. 1*H*). Such an alignment suggested that the branched plantlets are initiated sequentially inside the boat-shaped leaf. We confirmed this branch alignment by micro-computed tomography (CT) (Fig. 1*I*). This interpretation was also consistent with the observation that plantlets continuously protrude (see video at www.wolffiapond.net).

Where do branched plantlets arise? Focusing on the smallest primordium next to the inside surface of the boat-shaped leaf (Fig. 1 *J* and *K*), we observed several cells with a big nucleus and a condensed cytoplasm, arranged in the innermost region. Such cellular characteristics are typically associated with meristematic cells, suggesting that these cells may constitute the growth tip in *W. australiana*. Indeed, the development of the growth tip from branched plantlets supported this hypothesis: Following its initiation, the leaf primordium underwent asymmetric growth, with the outer portion of the primordium growing faster than the inner region. After the differentiation of the petiole (Fig. 1*J*), the fast-growing outer region protruded inward and overlapped with the slow-growing inner region (Fig. 1 *J* and *L*). The space between the two regions then formed a cavity inside the boat-like leaf, after which point a new growth tip initiated a new branch at the innermost point of the cavity (Fig. 1*L*). We thus conclude that the functional growth tip in *W. australiana* is not a multicellular cluster with a tunica-corpus structure like that seen in angiosperms, but only comprises one to a few cells that are induced during primordium differentiation. Such a growth tip organization repeated in each branch (compare Fig. 1 *M* and *J*).

While all branched plantlets grew outward, the innermost region next to the inner surface of the boat-shaped leaf enlarged upon flower induction conditions, visible as two bumps (Fig. 1 *N*–*P*). Compared to the specific orientation of floral organs (Fig. 1*Q*), these two bumps appeared to be the early stages of the stamen and the gynoecium primordia, respectively. Although we did not follow fertilization or seed development, we did observe the differentiation of floral organs (*SI Appendix*, Fig. S1; www.wolffiapond.net).

### Genome Sequence, Assembly, and Analysis

To aid in exploring the developmental innovations of the *Wolffia* genus, we performed whole-genome sequencing with a combination of strategies, including Nanopore PromethION (ultra-long), Illumina NovaSeq 6000 and Hi-C reads, and Bionano to construct the reference genome for accession wa7733 (referred here as Waus). We generated 18.46 Gb of ultra-long reads, 22.85 Gb of Illumina data, 253.08 Gb of Bionano data, and 36.55 Gb of Hi-C data (Dataset, Table S1). To help with later genome annotation, we produced 202.46 Gb of data by transcriptome deep sequencing (RNA-seq). To survey the genome, we used 11.3 Gb of Illumina paired-end reads (out of 22.85 Gb), resulting in 8.3 Gb of clean reads after quality-control steps and the removal of organellar and bacterial genomes. Based on these paired-end reads, we estimated the genome size to be 353,335,418 bp with a heterozygosity of less than 0.3% (*SI Appendix*, Fig. S2; Dataset, Table S2), which indicates that the Waus genome is very homozygous.

We used Nanopore reads to assemble the *W. australiana* genome into contigs. The G1 (genome version 1) contig length was 358.51 Mb, with an N50 size of 17.96 Mb. The G2 was 362.72 Mb (Dataset, Table S3). We also identified 4,345,907 bp of sequences corresponding to the mitochondrial and chloroplast genomes or to other contaminants. We distinguished true genomic sequences from other contaminants based on their GC contents (*SI Appendix*, Fig. S3 *A* and *B*). The Bionano data were used to correct the G3 genome, and five gaps were produced, resulting in 26 sequences for G4 (size of 358.77 Mb) that we assembled into 20 pseudo-chromosomes (Fig. 2 *A*–*C*; Dataset, Table S4). The mapping rates of RNA-seq against the genome assembly were 95.1% (Dataset, Table S5). Fluorescence *in situ* hybridization (FISH) analysis on prometaphase chromosomes confirmed 20 pairs of homologous chromosomes, representing 40 somatic chromosomes in the *W. australiana* genome (*SI Appendix*, Fig. S4).

**Fig. 2.**
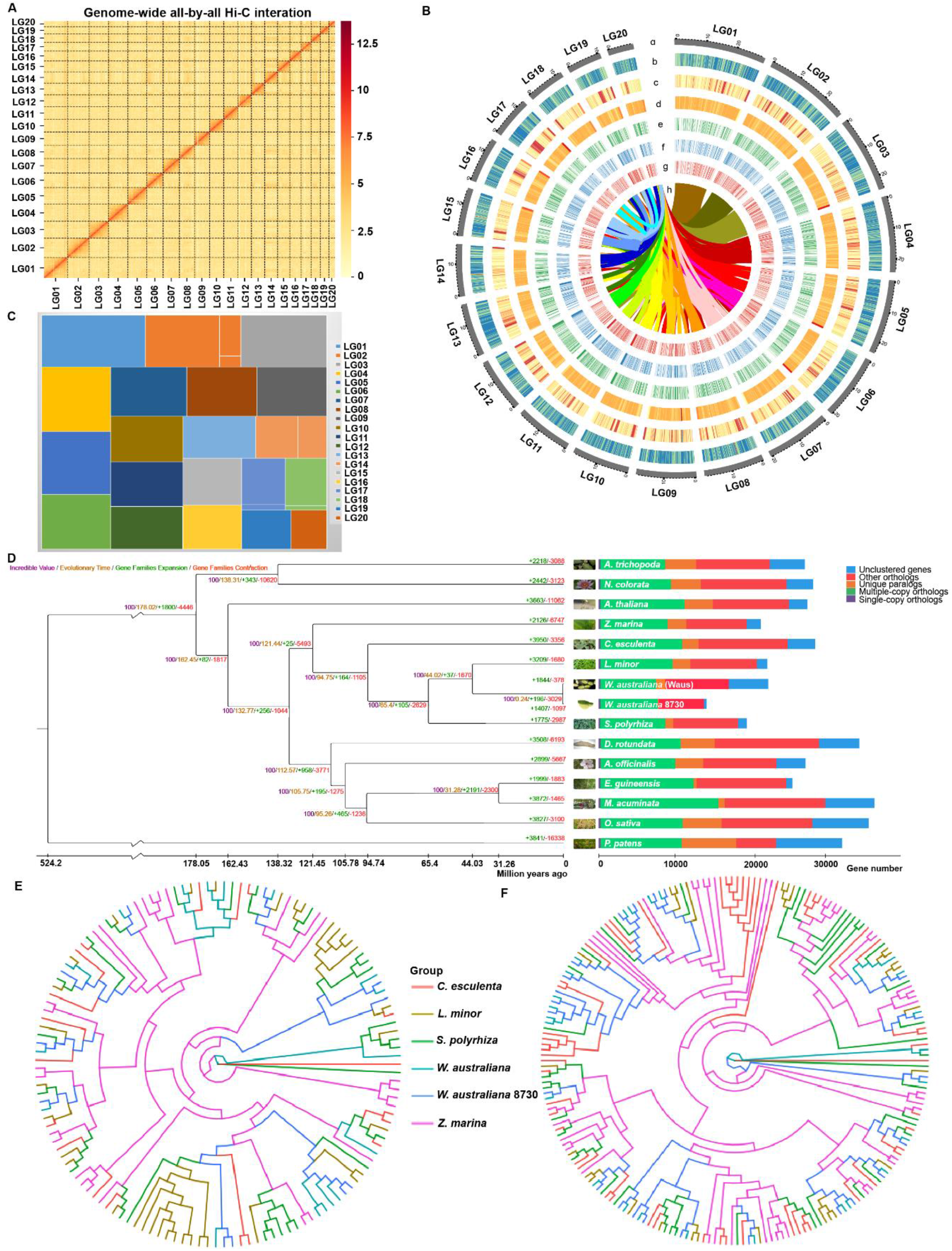
Genomic features of the *W. australiana* genome and gene family evolution in *W. australiana*. (*A*) Hi-C interaction Matrix for the 20 *W. australiana* pseudo-chromosomes. (*B*) Circos plot of the *W. australiana* genomic features: a, distribution of 20 chromosomes (each bar represents one chromosome, and the number represents the chromosome length); b, gene density; c, repeat sequence density; d, GC contents; e, gene density of Control transcriptome; f, gene density of Flowered transcriptome; g, gene density of Induced transcriptome. h, synteny and distribution of genomic regions across the *W. australiana* genome. (*C*) Treemap for contig length difference of 20 chromosomes. (*D*) Phylogenetic analysis of *W. australiana* and other plants. The single-cell green alga *Chlamydomonas reinhardtii* was used as outgroup. The value on each node represents the divergence time in millions of years (mya). Nodes marked red are published fossil calibration time points. Numbers marked in green/red represent expansion/contraction numbers on each branch. Photos on the right show the corresponding species. (*E*) *AGAMOUS-LIKE* (*AGL*) flowering-related genes. (*F*) Root-related *small auxin upregulated RNA* (*SAUR*) genes.

We also identified homozygous single nucleotide polymorphism (SNP) sites and insertion/deletion (InDels) by comparing the genome sequence reported here with those from other *W. australiana* species, yielding 6,764 SNPs and 3,166 InDels with minimal support of at least five Illumina reads (Dataset, Table S6). The contig N50 of our Waus genome was 18,579,918 bp compared to 251,357 bp for wa7733, 102,226 bp for wa8730, and 742,788 bp for wa8730 (three other published *W. australiana* genomes) (16, 17). Analysis of Benchmarking Universal Single-Copy Orthologs (BUSCO) demonstrated that our genome assembly is much more complete than previously sequenced and assembled *W. australiana* genomes, with BUSCO scores of 94.55% (Waus), 77.10% (wa7733), 80.29% (wa8730), and 87% (wa8730) (Table 1; Dataset, Table S7).

**Table 1.** Statistics of *W*.*a*. genome assembly comparison.

| Name | Waus | wa7733 (16) | wa8730 (16) | wa8730 (17) |
| --- | --- | --- | --- | --- |
| Total length (bp) | 358,772,296 | 359,766,217 | 337,899,876 | 456,810,926 |
| Contig number | 25 | 2,578 | 5,250 | 1,757 |
| Contig N50 (bp) | 18,161,740 | 256,298 | 102,418 | 734,533 |
| Longest contig (bp) | 28,936,433 | 1,664,978 | 679,034 | 7,663,058 |
| Scaffold number | 20 | 2,578 | 5,250 | 1,508 |
| Scaffold N50 (bp) | 18,579,918 | 836,551 | 109,493 | 1,169,370 |
| Longest scaffold (bp) | 28,936,433 | 5,333,369 | 1,714,878 | 8,358,235 |
| BUSCO | 94.55% | 77.10% | 80.29% | ~87% (?) |
| Mapping TGS | 99.50% | 98% | 96% | NA |
| Mapping NGS | 99.80% | 98% | 95% | NA |
| Genome size (Mb) | 358.8 | 359.8 | 337.9 | ~456. |
| Protein-coding genes | 22,484 | 15,312 | 14,324 | 22,293 |

### Genome Annotation and Phylogenetic Analysis

The *W. australiana* genome Waus contained 227.8 Mb (or 63.5% of the genome) of repetitive regions, consisting of both transposable elements (TEs) and tandem repeats (TRs). TEs with long terminal repeats (LTRs) represented the majority (41.8%) of TEs, followed by DNA transposons (9.2%), long interspersed nuclear elements (*LINE*s) (3.9%), and miniature inverted-repeat transposable elements (*MITE*s) (1.1%) (Dataset, Table S8). We predicted 1,658 non-coding RNAs for a total length of 237.2 kb, including 191 ribosomal RNAs (rRNAs), 867 non-coding RNAs (ncRNAs), eight regulatory RNAs, and 592 transfer RNAs (tRNAs) (Dataset, Table S9). We also predicted 22,484 protein-coding genes with an average length per gene of 3,789 bp (Dataset, Table S10 and S11). Finally, we compared the predicted Waus protein sequences to biological databases, resulting in functional annotation for 69.5% of all genes. We also obtained support for 75.04% of all predicted gene models in RNA-seq samples (Dataset, Table S10).

We determined the phylogenetic relationship between *W. australiana* and 14 other Viridiplantae species using a set of low-copy orthologous gene groups (Dataset, Table S11 and S12). Specifically, the phylogenetic results and fossil calibration time revealed that *W. australiana* 7733 diverged from *W. australiana* 8730 approximately 0.24 million years ago (mya) (Fig. 2*D*; Dataset, Table S13). We also compared the Waus genome to the genomes from other plant species to elucidate key genomic changes associated with adaptation to a small plant stature. We thus identified the expansion of 196 gene families and the contraction of 3,029 gene families in the cluster of *W. australiana* relative to plant species with larger body plans (Fig. 2*D*). For example, the *AGAMOUS-LIKE* (*AGL*) family involved in flowering typically clusters in 11 groups in flowering plants (Fig. 2*E*; Dataset, Table S14) but only defined nine groups in the Waus genome, suggestive of an incomplete *AGL* family in *W. australiana*. Similarly, many of the root-related small auxin up-regulated RNA (*SAUR*) genes were missing in *W. australiana* (Fig. 2*F*; Dataset, Table S14).

### Millifluidic Setup for Tracking Plantlets

To track the developmental processes of individual plantlets as they physically separate from one another, we designed a millifluidic chip system for plantlet culture. The chip design is quite simple: We poured polydimethylsiloxane (PDSM) (using Momentive clear RTV615 potting and encapsulating compound in a 10:1 ratio) into molds to create 1-mm-wide channels (Fig. 3*A*). Each chip can hold over 100 plantlets and is sufficient to carry out most experiments.

**Fig. 3.**
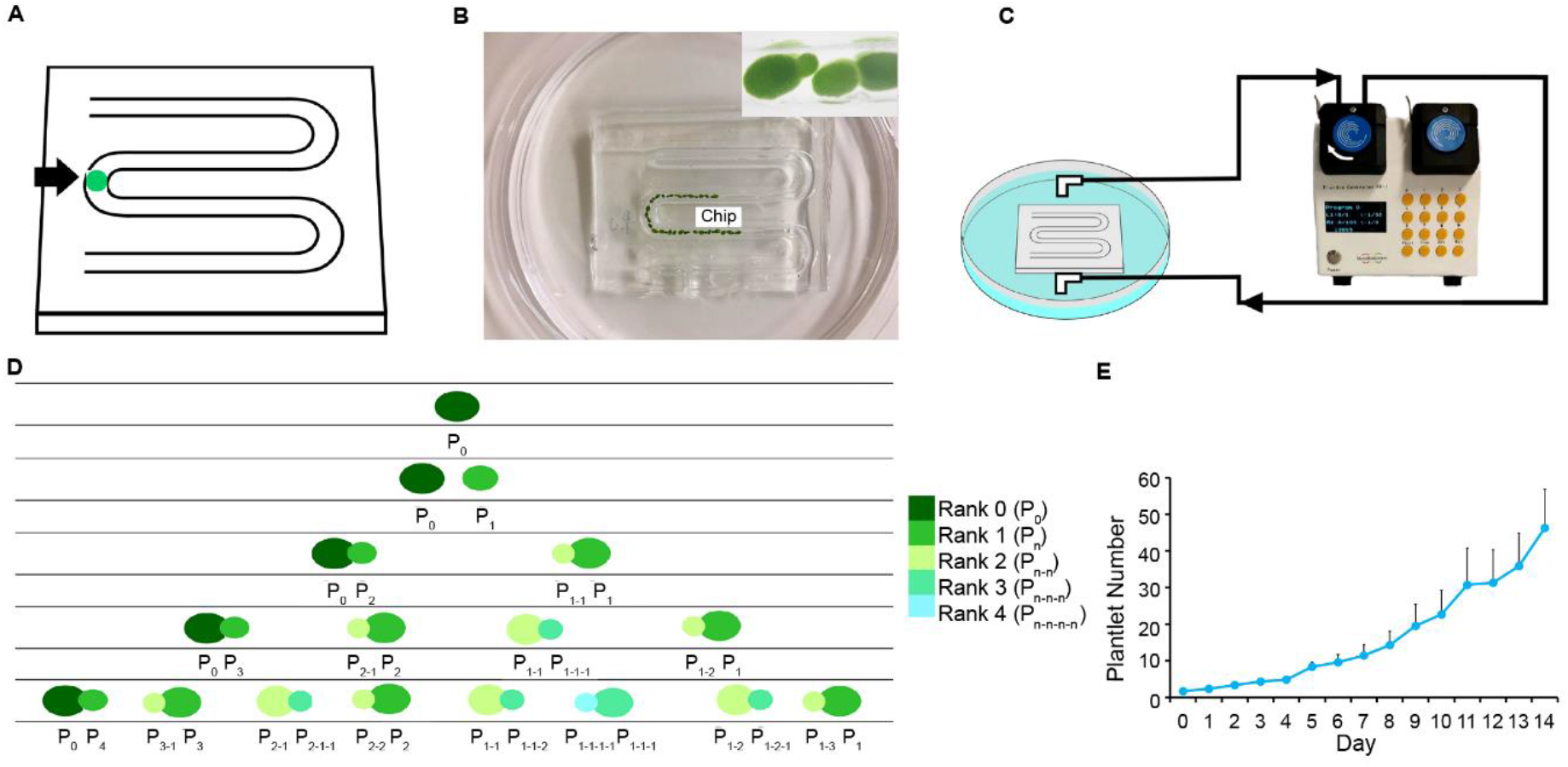
Plant-on-Chip culture platform. (*A*) Representative millifluidic chip (detailed information in Methods), showing a loaded plantlet. (*B*) Abscised plantlets (former branch) line up along the channel. (*C*) A peristaltic pump is connected to the chip to circulate liquid half-strength MS medium. (*D*) Diagram of growth pattern; the branch ranks are indicated by different colors. (*E*) Growth curve of cultured plantlets in the PoC in half-strength MS medium under SD conditions at 26 °C (n = 3).

To circulate the culture medium, we connected a peristaltic pump to the chip to help the liquid medium (half-strength Murashige and Skoog [MS]) flowing (Fig. 3*B*). Starting from a single plantlet with its first branch budding out along the long axis, the plantlets will line up along the channel (Fig. 3*C*). As the newly released (abscised) plantlet will also bud out of its own branch in the opposite orientation, the plantlets will align in a predictable order along the channel, as depicted in Fig. 3*D*. We designated the chip system “Plant-on-Chip” (or PoC).

For the convenience of tracking morphogenesis and manipulating the growth conditions, we also designed a customized incubator controlled by a computer and connected to a digital camera (*SI Appendix*, Fig. S5). All culture parameters can be programmed; plantlet growth can be monitored in an automated fashion. A typical growth curve under normal growth conditions (26°C, short-day photoperiod of 8 h light/16 h dark with 26.99 µmol photons/m^2^/s) is shown in Fig. 3*E*. Under normal conditions, each plantlet can release a new plantlet about every 48 h (±12 h) and survive for about a month thereafter. We tested several flowering inductive conditions previously reported to induce flowering in *W. microscopia* (7, 8, 18) and successfully defined conditions that will induce flowering in *W. australiana* growing within the PoC (*SI Appendix*, Fig. S6). We can therefore collect samples for morphological analyses (see above) and for the analysis of gene expression during flower development (see below).

### Genomic and Expression Characteristics Underlying Unique *W. Australiana* Morphological Traits

#### The Rootless Phenotype of W. australiana Cannot Be Explained by Gene Loss

A prominent morphological trait of *W. australiana* is its lack of roots. An attractive explanation would be that genes required for root development have been lost. However, we identified homologs for all known Arabidopsis genes involved in root development in the Waus genome (Dataset, Table S14). The formation of an auxin gradient is crucial for root initiation and maintenance of all root types (19, 20). However, in the region near the growth tip of *W. australiana*, cellular alignment required for an auxin gradient is hard to see (Fig. 1). This might result in the observed rootless phenotype.

#### Loss of NACs that Specify Vascular Differentiation may Explain the Lack of a Vasculature

We observed no vascular tissue in *W. australiana*, in contrast to its close relative, duckweed *Spirodela polyrhiza* (Fig. 4*A*). However, the cell wall composition of *W. australiana* is similar to that of aquatic plants harboring vascular bundles (21) (Fig. 4*B*). As xylem vessels usually possess thickened cell walls with specific patterns, referred to as the secondary cell wall (SCW), we looked for genes that are known to be involved in SCW formation in the Waus genome. We detected most SCW genes, indicating that cell wall biogenesis is likely intact in *W. australiana* (Dataset, Table S15). However, in contrast to the 13 Arabidopsis and nine rice SCW-related *NAM-ATAF1,2-CUC2* (*NAC*) gene family members (Dataset, Table S15 and S16), which encode the upstream master regulators for SCW formation in vascular plants (22), we only identified one homolog in *W. australiana*. This gene, WausLG14.977, clustered separately from the groups defined by Arabidopsis VASCULAR-RELATED NAC DOMAIN PROTEIN6 (VND6, At5g62380) and VND7 (At1g71930) (Fig. 4*C*), two master NACs involved in xylem vessel differentiation (22).

**Fig. 4.**
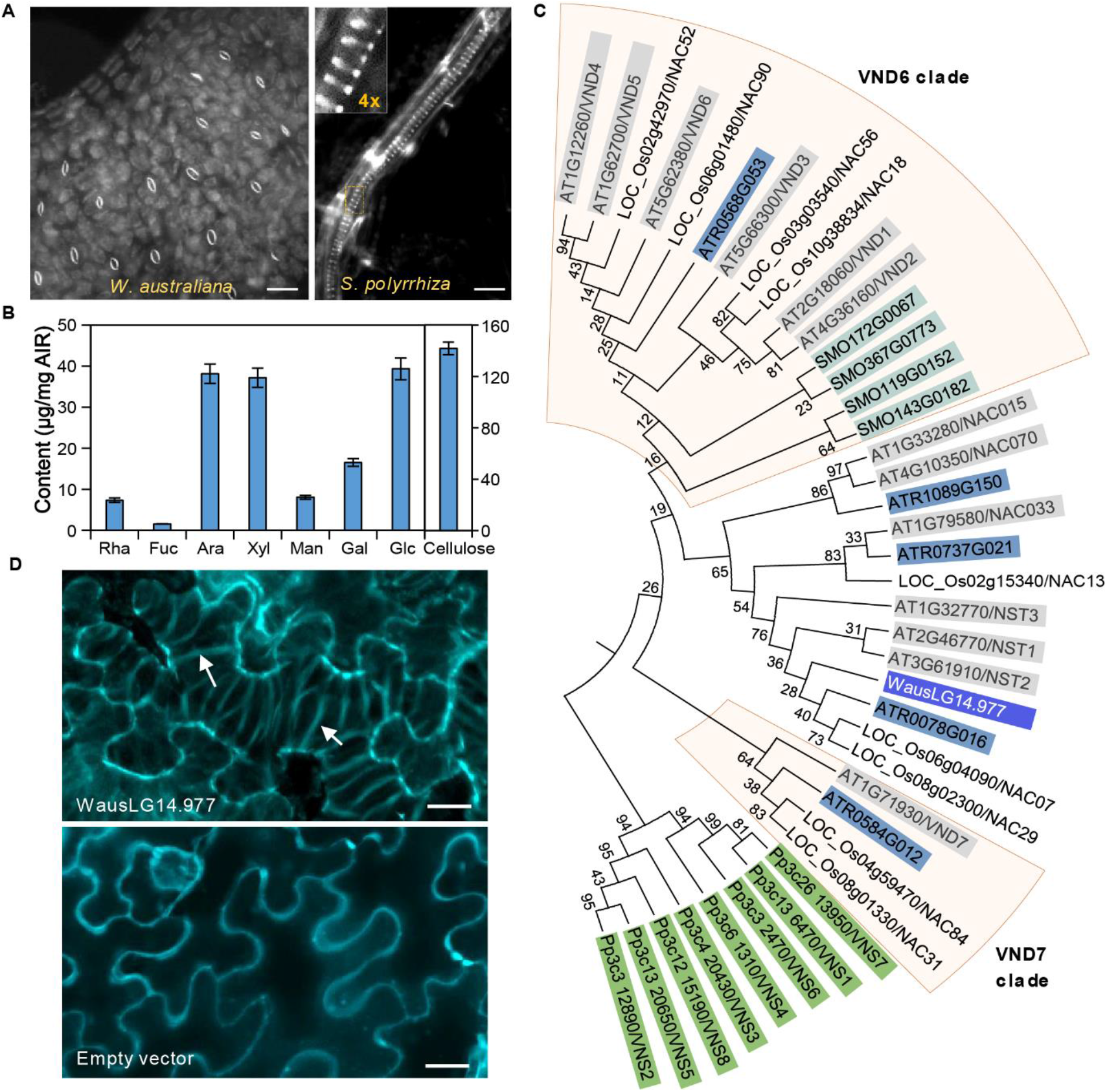
*W. australiana* lacks vasculature. (*A*) Staining of *W. australiana* and *S. polyrhiza* plantlets with the cell wall dye Direct Red 23, revealing no SCW vascular structure in *W. Australiana*, in contrast to the spiral-like xylem cells (inset) observed in the closely related duckweed *S. polyrhiza*. Bars = 80 μm (left) and 20 μm (right). (*B*) Cell wall composition of *W. australiana* plantlets. All components are shown with the scale to the left, except cellulose (right scale). Bar charts represent the mean ± standard deviation (SD) of five biological replicates. (*C*) Phylogenetic analysis of SCW-related NAC homologs in *W. australiana* and five representative genomes, indicating the absence of the VND homolog (boxed) in *W. australiana*. Pp, *Physcomitrium patens*; SMO, *Selaginella moellendorffii*; ATR, *Amborella trichopoda*; Os, *Oryza sativa*; AT, *Arabidopsis thaliana*. (*D*) Confocal images of *Nicotiana benthamiana* leaf epidermal cells transiently overexpressing WausLG14.977 or infiltrated with empty vector. Vessel-like cells were observed with WausLG14.977. Arrows indicate spiral SCW bands in a vessel-like cell. SCW, Secondary cell wall. Bars = 20 μm.

To test the function of WausLG14.977, we transiently expressed the gene in *Nicotiana benthamiana* leaves. Based on the UV excited fluorescent signals that derived from lignified SCW, we observed spiral SCW bands in the epidermal cells expressing WausLG14.977 but not in cells transiently infiltrated with the empty vector (Fig. 4*D*). The formation of vessel-like cells thus suggested that WausLG14.977 has the potential to be a functional VND-like regulator. This interpretation was further corroborated by the similarity in protein structure between the protein encoded by WausLG14.977 and Arabidopsis SECONDARY WALL-ASSOCIATED NAC DOMAIN PROTEIN1 (SND1, At1g32770), as predicted by RoseTTAFold (*SI Appendix*, Fig. S7).

Taken together, while the single *NAC* family member WausLG14.977 appeared to be functional for SCW formation, loss of homologs from the VND6 and VND7 clades and/or the lack of downstream components essential for xylem vessel development may be responsible for the absence of vascular tissues in *W. australiana*.

#### Correlation between Differential Gene Expression and Flower Development

Similar to *W. microscopia*, the transition from vegetative growth to induced flower morphogenesis is very fast and can take place in as few as 5 days in *W. australiana*, although not at a high frequency (*SI Appendix*, Fig. S6). This quick transition and the simple reproductive structures (Fig. 1), combined with the PoC system to track the transition process of a single plantlet, prompted us to explore the underlying regulatory mechanism. Accordingly, we collected three groups of individual plantlets for single plantlet RNA-seq: 1) plantlets with floral organs 5 days after application of EDTA treatment (to induce flowering), defined as F (flowered) samples; 2) plantlets with no floral organs under the same inductive conditions (plantlet responses to flowering induction were not synchronized), defined as I (induced) samples; and 3) plantlets grown for the same duration but not exposed to EDTA treatment, defined as C (control) samples (Fig. 5*A*). Using a candidate gene list of about 500 Arabidopsis and 100 rice flowering genes as queries, we identified about 200 homologous genes in *W. australiana* (Dataset, Table S14 and S16). Some of these genes exhibited differential expression between the C, I, and F groups, either upregulated (Fig. 5*B*; Dataset, Table S17) or downregulated (Fig. 5*C*; Dataset, Table S17). Among differentially expressed genes, we noticed that WausLG03.251 (homolog of *FLOWERING LOCUS T* [*FT*]) and WausLG11.346 (homolog of *FLOWERING PROMOTING FACTOR1* [*FPF1*]) are highly expressed in F samples (Dataset, Table S14 and S18), which was in agreement with the expression patterns of their Arabidopsis and rice homologs during the induction of flowering.

**Fig. 5.**
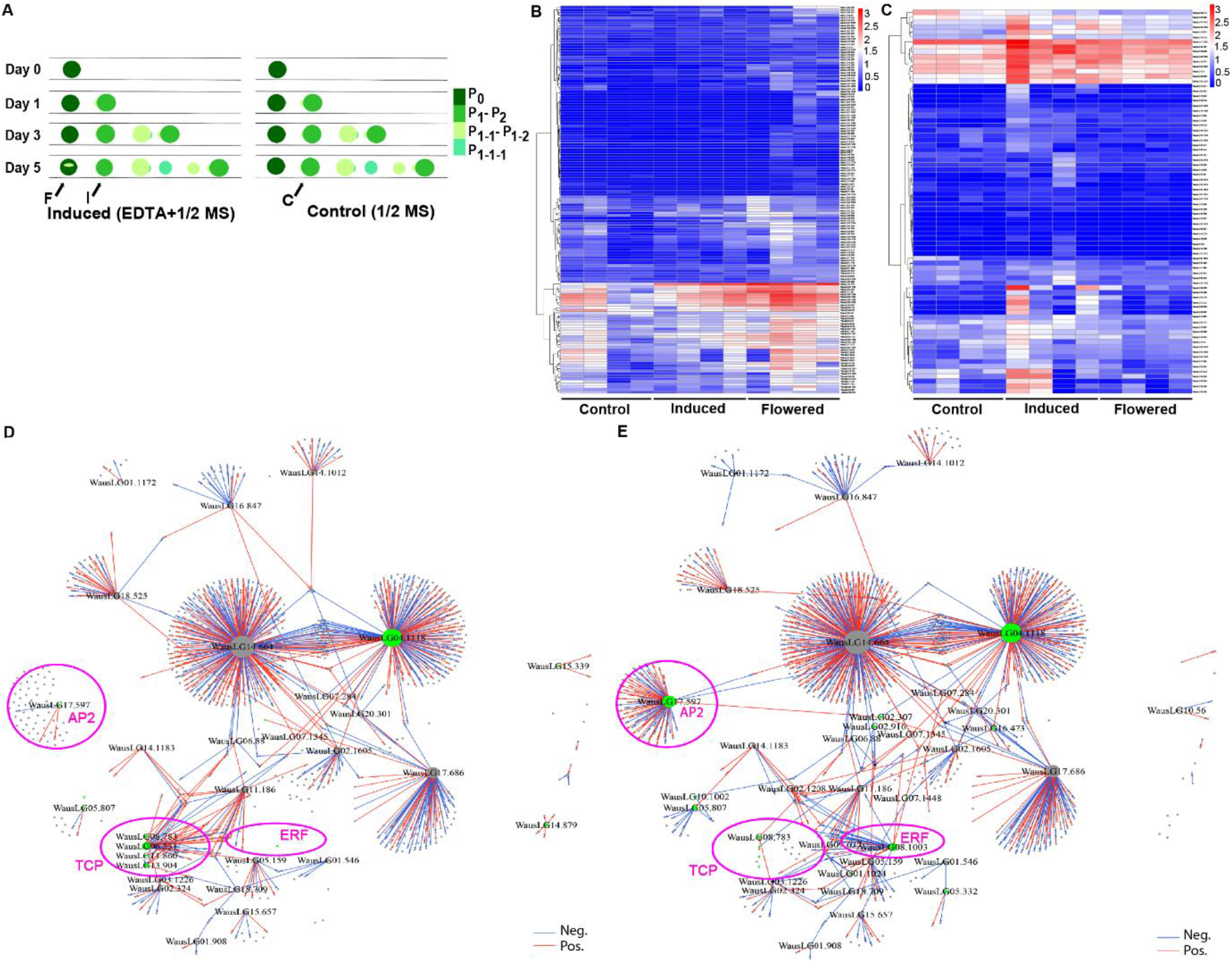
*W. australiana* floral induction and related transcriptome/transcription regulatory networks. (*A*) Diagram of the sampling design: F (flowered) samples were collected 5 days after culture under inductive conditions (left) from plantlets with a crack on the deck (shown as F with arrow). I (induced) samples were collected 5 days after culture under inductive conditions (left) from plantlets with no crack on the deck (shown as I with arrow). C (control) samples were collected 5 days after culture under non-inductive conditions (right, control) from plantlets (shown as C with arrow) remaining in a vegetative state. (*B*) Heatmap representation of 147 differentially expressed genes that are upregulated in Flowered compared to Induced (fold change ≥ 2 or ≤ –2, *P* ≤ 0.05). (*C*) Heatmap representation of 78 differentially expressed genes that are downregulated in Flowered compared to Induced (fold change ≥ 2 or ≤ –2, *P* ≤ 0.05). (*D*) TRN topological structure based on the comparison of the RNA-seq datasets from I and C samples. (*E*) TRN topological structure based on the comparison of the RNA-seq datasets from F and C samples. Red edges for positive regulation, blue edges for negative regulation, green nodes for differentially present genes, circled region for differentially present network in *D* and *E*. Pink circles highlight the nodes exhibiting topological differences between the two TRNs.

As an aquatic plant, the living conditions for *W. australiana* should be more stable than those of land plants because of the buffering capacity provided by water. Among the contracted gene families, we determined that miR156 is missing in *W. australiana* (Dataset, Table S14). Although its target transcripts from *SQUAMOSA PROMOTER-BINDING-LIKE PROTEIN3* (*SPL3*) or the other family members *SPL4, SPL5, SPL9*, and *SPL15* were present in the *W. australiana* genome, the loss of miR156, as well as miR172 (Dataset, Table S14), may explain the seemingly abrupt shift from vegetative growth to floral organ differentiation brought upon by the absence of the phase change observed in Arabidopsis and other plants normally mediated by the miR156-SPL module (23, 24).

Despite the loss of sepals and petals in the *W. australiana* flower, all MADS-box genes were present in the *W. australiana* genome and appeared to be expressed (Dataset, Table S14). This observation thus raises a question about the genomic maintenance of MADS genes participating in sepal and petal identity determination in an organism lacking these organs over the course of evolution.

We also carried out a transcription regulatory network (TRN) analysis (25) with single-plantlet RNA-seq data to decipher the regulatory mechanisms behind floral organ initiation and differentiation. While the overall topological structures of TRNs between C-I and C-F pairs were quite similar, several network nodes distinguished the two TRNs (circled in Fig. 5 *D* and *E*). While none of these distinct node genes were curated explicitly as known “flowering genes” (Dataset, Table S19), the node genes exhibiting the most prominent differences (*P* = 1.65e^−8^, odds ratio = 60.90, hypergeometric distribution test) between the two treatments were those annotated as encoding APETALA2 (AP2)/ETHYLENE RESPONSE FACTOR (ERF) and TCP (TEOSINE BRANCHED1 [TB1], CYCLOIDEA [CYC], (PROLIFERATING CELL NUCLEAR ANTIGEN FACTOR1 [PCF1]) members (Fig. 5 *D* and *E*; Dataset, Table S19; note: although WausLG06.531 and WausLG08.783 are circled, they do not belong to the TCP family). TCPs were reported to be involved in floral organ formation (26). However, we determined that none of the *TCP* genes are differentially expressed in the C-F TRN, as expected based on the proposed function of Arabidopsis *TCP*s, although they were differentially expressed in the C-I TRN, that is from samples induced to flower but lacking clear floral organs (Fig. 5 *D* and *E*). As AP2/ERF members have a wide range of functions in plants (26), the different TRN patterns for these two node genes raise interesting questions as to whether and how these genes function in *W. australiana* floral organ induction and differentiation.

### Single-nucleus RNA-seq, Protein Structure Predictions, and Gene Transformation

A valuable model system should have a high-quality sequenced genome and should be amenable to genetic manipulation for the dissection of gene function. Here, we applied single-nucleus RNA-seq (snRNA-seq) and other techniques with *W. australiana*.

In a pilot experiment, we collected 2 g of plantlets cultured in flasks under regular growth conditions for nuclear isolation (27), followed by snRNA-seq (28). After data processing and quality control, we retained 15,983 nuclei and 14,812 genes for clustering, cell type annotation, and other analyses. Fig. 6 illustrates the clustering pattern obtained by Uniform Manifold Approximation and Projection (UMAP); Dataset, Table S20 provides the cell type annotation of each cluster. We were surprised to identify almost all cell types (Dataset, Table S20; for full information see Dataset, Table S21), although the collected samples only consisted of boat-shaped leaves (with guard cells) and growth tips (Fig. 1). Notably, the results presented here are similar to those observed in snRNA-seq of freshwater sponge (*Spongilla lacustris*) (29). We will discuss this interesting phenomenon later.

**Fig. 6.**
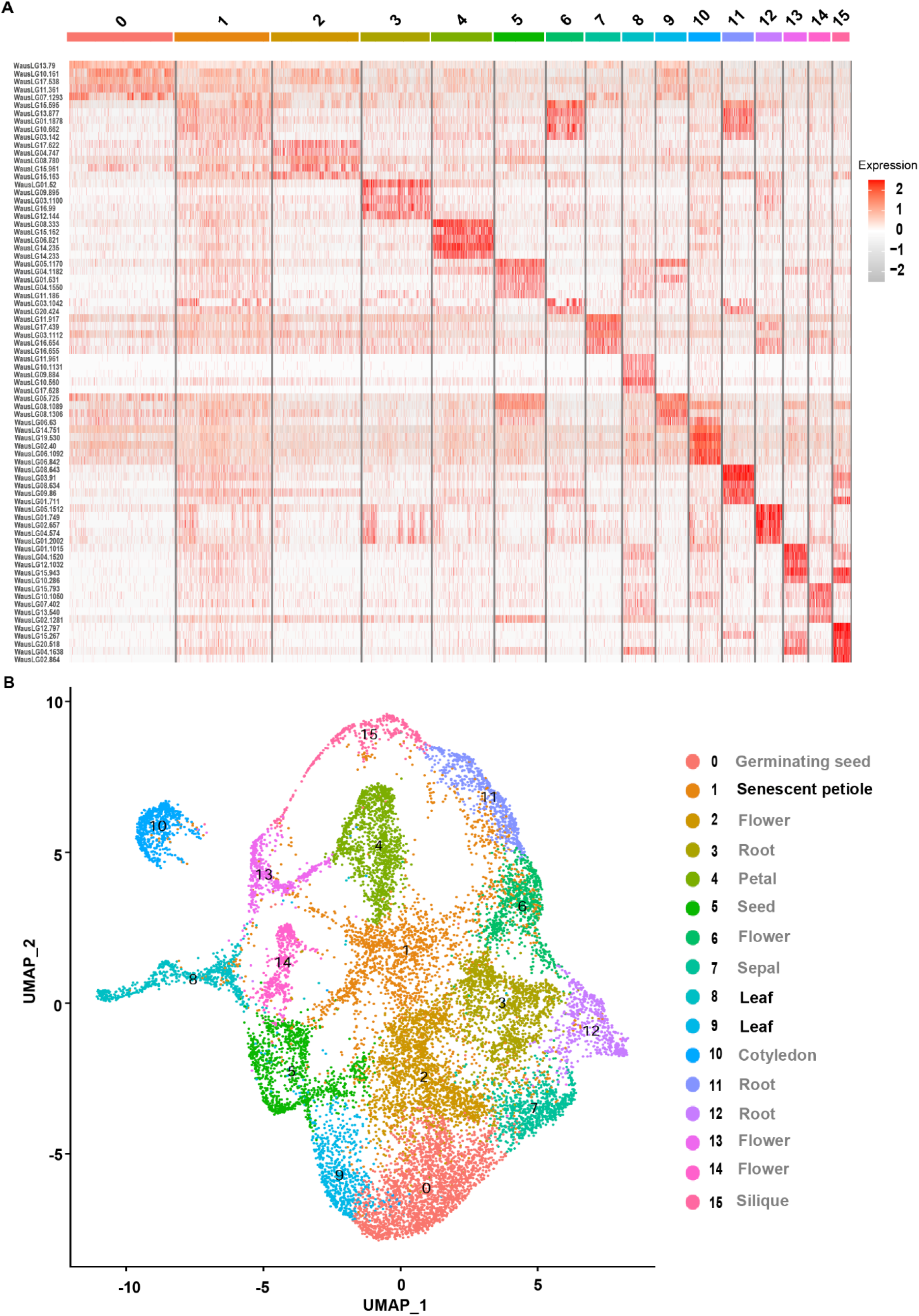
snRNA-seq reveals cell types absent in *W. australiana* plantlets. (*A*) Heatmap representation of differentially expressed genes across 15,983 cells clustered into 16 cell types. (*B*) UMAP visualization of 15,983 cells into 16 clusters (for detailed information, see Dataset, Table S16).

We also tested the usefulness of the protein prediction software AlphaFold2 (30, 31), considering the importance of protein structures in biological structures and processes. Accordingly, based on our high-quality genome sequence and RNA-seq data, we selected 6,800 predicted non-homologous protein sequences encoded by the *W. australiana* genome using MMseqs2 (32). With the non-docker version of AlphaFold2, we predicted 6,798 structures and related information (www.wolffiapond.net). Only two predicted proteins failed to generate a structure due to video memory limitations. While the true protein structures remain to be experimentally examined, the high efficiency of protein structure prediction with AlphaFold2 to the *W. australiana* coding sequences was very encouraging.

Effective genetic transformation is an indispensable tool to manipulate the genome during functional investigations. In duckweed research, transgenic procedures have been reported in species other than *W. australiana* (14). We previously developed transgenic procedures for the stable transformation of *Wolffia globosa* and *S. polyrhiza* (L.) (33, 34). Here, we identified a set of modifications to the procedure to generate transgenic *W. australiana* plants (*SI Appendix*, Fig. S8).

The PoC system is available for research teams interested in investigating fundamental questions in plant biology and plant development.

## Discussion

Here, we report that using *W. australiana* as a model system enables some fundamental issues in plant morphogenesis to be analyzed.

### Uncoupling the Form and Function of the Growth Tip in Angiosperms

The growth tips of most angiosperms exhibit a tunica-corpus structure, while those of gymnosperms, pteridophytes, and bryophytes do not. Since the growth tips nevertheless carry out their functions as the centers of morphogenesis, the functional relevance of the tunica-corpus structure in angiosperms and how multicellular growth tips emerged remain elusive. Based on our observations in *W. australiana*, it is clear that the cell(s) located in the innermost regions of the cave behave as a growth tip, although they are not organized into a tunica-corpus structure (Fig. 1*K*). Under non-inductive conditions, the growth tip continuously generates leaf primordia (Fig. 1; *SI Appendix*, Fig. S2). Under inductive conditions, the growth tip enlarges and protrudes further inward before forming one stamen and one gynoecium (Fig. 1 *N***–***Q*). Based on these observations, we conclude that the tunica-corpus structure is dispensable for the proper function of the angiosperm growth tip, at least in the case of *W. australiana*.

The uncoupling of the form and function of the growth tip in angiosperms is not exclusive to *W. australiana*. Indeed, mutants in the Arabidopsis gene *WUSCHEL* (*WUS*) retain the ability to produce lateral organs from their growth tip for flowering, although the mutant lacks a typical tunica-corpus structure (35). One outstanding question worth pursuing is to investigate when and how the tunica-corpus structure evolved at the growth tip of angiosperms.

The emergence of axillary meristems has been explained by two rival hypotheses: *de novo*, the meristematic cells arise from differentiated somatic cells; and detached, the meristematic cells derives from preexisting meristematic cells (36). In agreement with a previous report (37), our observations on growth tip emergence during the differentiation of leaf primordia (Fig. 1 *I, J, L* and *M*) support the *de novo* hypothesis. In addition, compared to the *in vitro* regeneration of a shoot apical meristem in tissue culture conditions (38), the relatively stable pattern of growth tip emergence in *W. australiana* makes it as an ideal experimental system to precisely investigate the spatiotemporal mechanism of the transition from partially differentiated somatic cells to meristematic cells.

### A Unique Opportunity for Detailed Analyses of the Causal Relationship between Genome Rewiring and Morphogenetic Simplification

The genome of *W. australiana* did not exhibit a dramatic size reduction relative to its close relatives *Colocasia esculenta* and *Zostera marina*, which produce multiple organ types in contrast to *W. australiana* (Fig. 2). Based on the genomic analyses presented here, we offer several clues that might explain this morphogenetic simplification: First, the *W. australiana* genome might have experienced a dramatic structural rewiring, as we detected 378 expanded gene families and 1,844 contracted gene families based on gene copy number, relative to its closest relatives (Fig. 2). Second, specific gene families showed a contraction in their constituent members. Members from the *AGL* family cluster into 11 groups in most plant species, whereas the *W. australiana* genome encoded AGL homologs belonging to nine groups (Fig. 2*E*). Similarly, the 13 Arabidopsis SCW-related *NAC* genes had a single homologous gene in *W. australiana*, which might be responsible for the non-vascular phenotype of this tiny plant (Fig. 4, Dataset, Table S15). Based on recent findings of the dynamic features of chromatin structures, it is reasonable to hypothesize that the relative stable aquatic conditions experienced by *W. australiana* may have contributed to genome rewiring, including gene and/or gene family contraction.

Other regulatory mechanisms may have also participated in the morphogenetic simplification of *W. australiana*. For example, the simple absence of a gene or gene family cannot explain all cases of morphological innovations, as we identified almost all known genes required for Arabidopsis root development in the *W. australiana* genome (Dataset, Table S14). More curiously, our pilot snRNA-seq experiment revealed the expression of genes that are typically markers for tissues or organs missing in *W. australiana* plantlets. Notably, the genome of freshwater sponges was recently shown to harbor genes involved in nerve cell development, although this organism lacks this cell type (29). It is possible that the expression of certain genes may occur prior to the emergence of a given morphogenetic event. Before the genes were coopted for specific morphogenetic events, they may have carried out other functions. Our high-quality genome opens the door to exploring the relationships between gene annotation and morphogenetic events.

### Retracing Cell Fate Transition from Vegetative to Reproductive Growth in an Individual Growth Tip

Flowering is one of the morphogenetic processes in angiosperms that has attracted the most interest in plant research. While significant progress has been made over the past century, a thorough dissection of the underlying regulatory mechanisms has been hindered by two limitations: 1) the transition from vegetative to reproductive growth is a long process consisting of sequential changes from photosynthetic leaves to peripheral organs, with the added interference of complex branching in an asynchronous system; and 2) the transition from vegetative to reproductive growth integrates multiple internal and external environmental changes within each growth tip, which may respond quantitatively and qualitatively differently (2). A simpler experimental system would be especially helpful here.

With the PoC system described here, we easily traced the entire transition from vegetative to reproductive fate of an individual growth tip. Furthermore, we were able to trace how the growth tip differentiates into two primordia upon flowering induction (Fig. 1 *N***–***P*), which will further differentiate into the stamen and gynoecium, all within a time frame of a few days (Fig. 1 *Q, F* and *G*). In addition, we identified simple but efficient inductive conditions to trigger the transition. Therefore, with the assistance of cutting-edge single-cell techniques, this PoC system should provide a unique opportunity to decipher the regulatory mechanisms that drive fate transition with a much lower signal-to-noise ratio.

The simplicity of the *W. australiana* floral structure and accessibility may offer a novel approach to answering Darwin’s abominable mystery: the origin of the flower. Based on our observations (Fig. 1 *N*–*Q*), the stamen and gynoecium are aligned at the ends of separated branches. If the stamen and gynoecium in *W. australiana* are considered as elaborated heterosporangia (39, 40), the origin of the flower may be seen as a result of two separate evolutionary innovations: the transition from homosporangium to heterosporangia and the combination of individually initiated fertile telomes, each with micro-and mega-sporangium, together as a morphological unit recognized as a “flower.” *W. australiana* may thus be amenable to the exploration of how the two primordia emerge upon induction and how they differentiate into a stamen or gynoecium. Such dissection should provide a substantial basis for the investigation of heterosporangia differentiation in more species from Pteridophytes to Spermatophytes.

Although we have yet to explore fertilization and seed formation in the *W. australiana* life cycle, we did precisely visualize and trace most of the morphogenetic events required for sporophyte development at the cellular level. We have illustrated the possible causal relationship between morphological traits and genome variation for root and vascular differentiation. Other interesting morphogenetic events will be open to investigation, such as the programmed cell death that results in the formation of the crack on the “deck” of the boat-like plantlets; how guard cells differentiate on the “deck” but not on the “hull”; and how the asymmetric growth of the leaf primordium is guided. The PoC system established here can offer a unique opportunity for deciphering the regulatory mechanisms of core processes of plant development and other interesting morphogenetic events during the sporophyte stage.

## Materials and Methods

### Biological materials

*Wolffia australiana* wa7733 was from Prof. Hongwei Hou Lab at the National Aquatic Biological Resource Center, Institute of Hydrobiology, Chinese Academy of Sciences. *W. australiana* plants used for genome sequencing and transcriptome sequencing were cultured in half MS (Solarbio) solid medium (1% sucrose, pH 5.5) under long-day (LD: 16-h light/8-h dark) conditions at 26 °C. The fresh plants cultured for 10-15 days were used for the extraction of genomic DNA for genome sequencing and total RNA for transcriptome sequencing from plants in a population under cultured conditions (Dataset, Table S1).

The *Wolffia* plants were grown in liquid half MS medium (1% sucrose, pH 7 and antibiotics free), under short-day (SD: 8-h light/16-h dark) conditions at 26 °C on the plates for 1-2 weeks. Plants were transferred to half MS medium (Control, C) or half MS + EDTA (Sigma) medium (Induced but not flowered, I and Flowered, F) in chip for 5 days before RNA extraction for single-plant RNA-seq. Each group (C, I, F) included four single *Wolffia* plants as four biological replicates.

The *Wolffia* plants were grown in half MS medium, under short day at 26 °C on the plates. The fresh weight of *Wolffia* was 2 g for one sequencing for single-nucleus RNA-seq.

### Additional Methods

Floral organ staining, Photomicrograph conditions, Plant-on-chip device design, Cryo Scanning Electron Microscopy (Cryo-SEM), Sample preparation for Micro Computed Tomography (microCT) and Transmission Electron Microscopy (TEM), Genome sequencing, Genome size estimation, Chromosome-level genome assembly and assessment, Genome repeat and gene annotation, Genome phylogenetic analysis, RNA extraction and library preparation for single-plant RNA sequencing, Analysis of genes related to morphological processes, Differential expression genes (DEGs) analysis, Chromosome FISH analysis, Microscopy, Cell wall composition analysis, Phylogenetic analysis, Transcription regulatory network (TRN) analysis, Protein structure prediction and classification, Single nucleus isolation, single-nucleus library construction and sequencing, raw data processing, and generation of gene expression matrix, Cell clustering, cell type identification, quality control and cell type annotation for snRNA-seq, Gene transformation of *W. auatraliana*, and Statistical Analysis are in *SI Appendix*, SI Materials and Methods.

## Supporting information

Supplementary Tables with dataset.

## Data availability

The genome sequence of *Wolffia australiana* wa7733 has been deposited at NCBI Genome under the accession CP092600-CP092619 for 20 chromosomes. Raw genome and transcriptome sequencing reads have been deposited into the NCBI sequence read archive (SRA) under the BioProject PRJNA808652 (for Nanopore), PRJNA808655 (for Illumina genome), PRJNA808685 (for Hi-C), PRJNA808734 (for BioNano), PRJNA808736 (for RNA-seq for genome), PRJNA808739 (for single-plant RNA-seq), and PRJNA809022 (for single-nucleus RNA-seq). BioNano data have also been deposited into NCBI Supplementary Files under the accession SUPPF_0000004267. Genome sequence, single-plant and single-nucleus RNA-seq are also available at the *Wolffia australiana* wa7733 genome database: http://wolffiapond.net.

## ACKNOWLEDGMENTS

We are grateful to Hai-Qi Meng, Mei-Qin Chen in the Bai lab at Peking University (PKU) for pilot experiments with *Wolffia globosa*; Professor T. Oyama at Department of Botany Kyoto University for providing *W. australiana* 7733; Yi-Zhen Jia, Xiao-Fan Zheng, Yue-Yi Che and others in Dr. Feng Li’s Lab at High School Affiliated to Renmin University (HSARU) for their participation in establishing the PoC system and pilot morphological analysis; Xing Zhang of HSARU for designing the cover; Kai-Le Wang for manufacturing the incubator; Shu-Qiang Chen and Hui Zhang from BGI-Shenzhen for help with snRNA-seq data; Prof. Wen-Qin Wang of Shanghai Jiaotong University for providing information about duckweed genome studies; Prof. Yue-Hui He of the School of Advanced Agriculture, PKU, Prof. Yun-Yuan Xu and Jing-Yu Zhang of the Institute of Botany, Chinese Academy of Sciences, Prof. Tong-Da Xu of Fujian Agriculture and Forest University, Prof. Hong-Chang Cui of Florida University for the Arabidopsis gene lists involved in flowering, auxin and root development. We thank the Facilities Cores at National Center for Protein Sciences and the TEM platforms at the College of Life Sciences of Peking University for cryo-SEM, TEM and micro-CT. We also thank Jianlin Chen at the College of Engineering of Peking University for cryo-SEM, TEM and micro-CT. This work was crowd-funded by all the authors. This work was supported by Mississippi State University (to L.L.), the International Partnership Program of Chinese Academy of Sciences (152342KYSB20200021) (to H.-W.H.), the National Natural Science Foundation of China (32001107) (to J.-J.Y.), and the National Natural Science Foundation of China (10721403) (to Q.O.).

## Notes

### Competing Interest Statement

The authors have declared no competing interest.

