## Supplementary Materials and Methods, Figures and Table. for "Plant-on-Chip: core morphogenesis processes in the tiny plant *Wolffia australiana*"

This file includes:
SI Materials and Methods
Figs. S1 to S8
Table S-Materials
SI References

### SI Materials and Methods

**Floral Organ Staining.** The flowered *Wolffia australiana* was (i) stained in Direct Red 23 (1/100) over 8 h; (ii) transferred to transparent solution (6 g trichloroacetaldehyde in 0.6 mL glycerol and 2.2 mL $H_2O$) until clear; (iii) washed in $H_2O$ 3 times with gentle shaking.

**Photomicrograph Conditions.** We photographed and shot time-lapse recording with a microscope digital camera (Olympus, DP27) or incubator's camera. The shooting time was based on the experimental design.

**Plant-on-chip Device Design.** We developed a light culture method to study *Wolffia* using a new plant-on-chip system. The chip was made of PDSM (mixed Momentive and RTV615). PDSM (the ratio of Momentive:RTV615 is 10:1) was poured into mold. The mold was placed into a 70 °C oven for 4 h, and cut into pieces before use. The mold was printed by a 3D printer (*SI Appendix*, Fig. S5), with 1-mm channel width, 10-mm depth, and 35-mm length. A single *W. australiana* was planted into the channel of chip in clean bench and submerged in medium. The set was connected to the pump (MesoBioSystem) for medium cycling (the pump rate could be set 0-8 times/sec). All the solutions and components were sterilized before use because there were no antibiotics. The set and pump were transferred into incubator without destabilization. This custom-made incubator includes 5 systems: (1) light, (2) thermostat, (3) camera and microscope, (4) control and display, and (5) cabin (*SI Appendix*, Fig. S5). The blueprint for the incubator was shown in *SI Appendix*, Fig. S5. The culture conditions could be set on the touch-screen controller or the external computer.

**Cryo Scanning Electron Microscopy (Cryo-SEM).** Cryo-SEM was used to study the *Wolffia* morphology. The equipment included the Helios NanoLab G3 UC scanning electron microscope (Thermo Fisher Scientific) and the PP3010T workstation (Quorum Technologies), which had a cryo preparation chamber connected directly to the microscope. The *Wolffia* were frozen in subcooled liquid nitrogen (-210 °C) and transferred in vacuum to the cold stage of the chamber, where sublimation (-90 °C, 5 min) and sputter coating (10 mA, 60 s) with platinum were conducted. Finally, the samples were transferred to another cold stage in the scanning electron microscope and imaged. The image was recorded using the electron beam at 5 kV and 0.2 nA with a working distance of 10 mm. The resolution of the final data was 3072 x 2048.

**Sample preparation for Micro Computed Tomography (microCT) and Transmission Electron Microscopy (TEM).** *Wolffia* plantlets were fixed with 2% (w/v) glutaraldehyde (Sigma-Aldrich, G7651) and 2% (w/v) paraformaldehyde (Sigma-Aldrich, P6148), 0.1 M phosphate buffer (81 mM $Na_2HPO_4$ (Sigma-Aldrich 71649) and 19 mM $NaH_2PO_4$ (Sigma-Aldrich 71507)), pH 7.4, for 2 h at the room temperature, and then held overnight at 4 °C. The next day, the samples were washed with 0.1 M phosphate buffer for three times and post-fixed in 2% (w/v) $OsO_4$ (Ted Pella, 18459) and 1.5% (w/v) potassium ferrocyanide (Sigma-Aldrich, P3289) for 2 h at 4 °C. Following five washes with phosphate buffer and distilled $H_2O$, samples were put into 1% thiocarbohydrazide (Sigma-Aldrich, 223220) and followed by another post-fixation in 1% (w/v) $OsO_4$, and three times of washing with distilled $H_2O$ were also included after each step. At last, samples were dehydrated through a graded alcohol series and embedded in Spurr's resin (SPI Supplies, 02680-AB).

Resin blocks were used for microCT and the pictures were taken by Xradia Context (Zeiss) or SkyScan 1272 (Bruker). Next, selected resin blocks were cut into 70-nm sections using an ultramicrotome (Leica Microsystem, UC7) referring to the microCT data for TEM (JEOL, JEM-1400).

**Genome Sequencing.** Genomic DNA was extracted from a *W. australiana* sample collected from a population using SDS method to construct ultra-long DNA libraries using above 10 μg DNA (with size selected ≥ 50 kb) and the SQK-LSK109 sequencing preparation kit (Nanopore, Ligation Sequencing Kit). The samples were sequenced on the Promethion (Oxford Nanopore Technologies, UK) at the Genome Center of Grandomics (Wuhan, China). The same genomic DNA was used to produce the short-read Illumina sequencing library by performing g-TUBE fragmentation, repair, adaptor connection, PCR amplification and recycling approximately 400-bp sequence using approximately 2 μg DNA and then sequenced on the Illumina NovaSeq 6000 platform at Grandomics (Wuhan, China). As for the Hi-C data, freshly prepared samples were chopped into 2-cm pieces and vacuum infiltrated in nuclei isolation buffer supplemented with 2% formaldehyde. Crosslinking was stopped by adding glycine and additional vacuum infiltration. Fixed tissue was then grounded to powder before re-suspending in nuclei isolation buffer to obtain a suspension of nuclei. The purified nuclei were digested with 100 units of DpnII and labelled by incubating with biotin-14-dCTP. Biotin-14-dCTP from non-ligated DNA ends was removed because of the exonuclease activity of T4 DNA polymerase. The ligated DNA was sheared into 400-bp fragments, and then was blunt-end repaired and A-tailed, followed by purification through biotin-streptavidin-mediated pull down. Next, for Bionano physical mapping, DNA extracted from population plants of *W. australiana* were subject to library preparation with Bionano Prep$^{TM}$ Plant DNA Isolation Kit (Bionano Genomics, Cat# 80003) following manufacturer recommended protocols. Optical scanning was provided by Bionano Genomics (https://bionanogenomics.com), with Bionano Prep DLS Labeling DNA Kit (Bionano Genomics, Cat# 80005). Labelled DNA samples were loaded and run on the Saphyr system (Bionano Genomics) in Grandomics. Finally, the Hi-C libraries were quantified and sequenced using the Illumina NovaSeq 6000 platform. To obtain RNA-seq, total RNA was extracted from different samples in TRIzol reagent (Invitrogen, Cat# 15596018)/CTAB-LiCl method (Plant) on dry ice and processed following the manufacturer's protocol. Sequencing libraries were generated using TruSeq RNA Library Preparation Kit (Illumina, USA) following standard protocol. Briefly, about 1 μg RNA per sample was used and enriched from total RNA using oligo(dT)-attached magnetic beads. The first-strand cDNA was synthesized with random primer and M-MLV Reverse Transcriptase, and the second-strand cDNA synthesis was followed by using DNA Polymerase I and RNase H. The synthesized cDNA was end-repaired, A-tailed and ligated to the sequencing adapters according to library construction protocol. The cDNA fragments were selected by AMPure XP beads (Beckman Coulter, USA) to an average size of 150-200 bp and amplified by PCR with Phusion High-Fidelity DNA polymerase, Universal PCR primers and Index Primer. At last, PCR products were purified with AMPure XP Beads (Beckman Coulter, USA) and library quality was assessed on the Agilent Bioanalyzer 2100 system. After that, the library was sequenced on the Illumina NovaSeq 6000 platform.

**Genome Size Estimation.** The genome size of *W. australiana* was estimated using the K-mer method based on the Illumina NovaSeq 6000 next-generation sequencing (NGS) data. Firstly, the quality-filtered paired-end data were filtered by Fastp v0.19.4 with specific parameters (-f 3 -t 2 -F 3 -T 2 -n 0) and then sequences of bacteria, mitochondria and chloroplast were removed from the quality-filtered reads to produce the complete genome data by aligning against a bacteria and organelle database with BLASTN v2.9 with the parameter "E-value $1e^{-5}$".

Secondly, the K-mer size was selected to be 17-bp and total number and average depth of K-mer sequences were calculated from the complete genome data reads (Dataset, Table S2). Finally, the genome size was obtained by calculation based on the formula: genome size = total K-mer number / average K-mer depth.

**Chromosome-level Genome Assembly and Assessment.** NextDenovo v2.0-beta.1 (https://github.com/Nextomics/NextDenovo/) was used with the filtered ultra-long sequences to produce the initial genome sequence (G1) with these parameters "reads_cutoff = 1k seed_cutoff = 50k" and "-n 1966 -q 0 -i 0.41 -s 0.18 -n 2 -r 0.14 -m 7.88 -c 80 -z 12". Then the ultra-long passed reads and the genome quality-filtered paired-end reads were applied to polish the G1 genome sequences by NextPolish v1.0.5 with the parameters (task = 55512121212) to produce polished genome (G2). To eliminate contamination sequences of the genome which may result in potential problems in the downstream analysis, the G2 contigs were aligned to the Nucleotide Sequence Database (NT) by using BLASTN v2.9 with the parameter "E-value 1e-5" and the sequence alignment results were classified based on species taxonomy. The contig was aligned to bacteria genome and was classified to contamination sequences and filtered out from G2 to form the decontaminated genome (G3).

To evaluate the accuracy of the G3, Illumina genomic pair-end reads were mapped to the genome contig sequences by the "mem" submodule of BWA. The mapping identity and genome coverage of the genome assembly were calculated based on the mapping result obtained by SAMtools v1.4 with default parameters. Homozygous single-base variations were subsequently detected by using BCFtools v1.8.0 with default parameters. Furthermore,

Illumina RNA-seq reads were mapped to the genome sequence by using HISAT2 v2.1 with default parameters and the mapping rate of RNA-seq reads was calculated by SAMtools. The completeness of conserved genes and eukaryote core genes assembly were evaluated by using BUSCO v3.1.0 with the 'embryophyta_odb10' dataset and CEGMA v2 with default parameters, respectively. Meanwhile, 10-kb bins were formed from G2 and were used to build the distribution of GC-Depth.

To obtain accurate genome, the Bionano physical mapping was used to map the G3 by Bionano Solve™ data analysis software to form the G4 version of genome. To anchor the G4 sequences to 20 chromosomes, the original Hi-C paired-end reads generated by the Illumina platform were aligned to the final contig sequences by Bowtie2 v2.3.2 with parameters "-end-to-end --very-sensitive -L 30". The contigs were clustered and ordered by using LACHESIS (41) with parameters "CLUSTER MIN RE SITES = 100; CLUSTER MAX LINK DENSITY = 2.5; CLUSTER NONINFORMATIVE RATIO = 1.4; ORDER MIN N RES IN TRUNK = 60; ORDER MIN N RES IN SHREDS = 60". Finally, the genome contigs were scaffolded into chromosome sequences to get the chromosome-level genome (G5) sequences (Dataset, Table S3) and Hi-C interaction heatmap was obtained based on the contig interaction results.

**Genome Repeat and Gene Annotation.** Simple sequence repeats (SSR) and tandem repeat sequences (TR) were firstly identified from the chromosome-level genome by GMATA v2.2 with default parameters and Tandem Repeats Finder v4.07b with parameters "2 7 7 80 10 50 500 -f -d -h -r", respectively. Sequence identified as tandem repeat sequences were 'soft masked' as lowercase letters in the genome. Secondly, long terminal repeat (LTR) retrotransposons were identified from the TR-masked genome assembly using LTR_finder v1.07 and LTR_harverst v1.5.10 with default parameters separately; a LTR-repeat library was constructed by LTR_retriver v1.8.0 with default parameters based on the results above. Miniature inverted transposable elements (MITEs) were subsequently identified using MITE-Hunter v11-2011 (42) with parameter "-n 20 -P 0.2 -c 3". The identified LTRs and MITEs were combined to mask the *W. australiana* genome and novel transposable elements (TEs) were identified by RepeatModeler v1.0.11 (https://github.com/Dfam-consortium/RepeatModeler) to construct the TE library. Finally, LTRs library, TEs library and Repbase were combined into one library file as the input for RepeatMasker with parameters "nolow -no_is -gff -norna -engine abblast -lib lib" to search repetitive sequences throughout the genome.

Protein-coding genes were predicted from the genome using the EVidenceModeler (EVM) pipeline v1.1.1 which integrates gene models originated from three sources of predictions: De novo, protein homology and transcriptome prediction. Homologous proteins of 15 plant species (Dataset, Table S11 and S12) across the Viridiplantae were used by GeMoMa v1.6.1 (43) with default parameters. Illumina RNA-seq reads were applied to predict genes by using HISAT2 (default parameters), StringTie v1.3.3d (default parameters) and PASA v2.3.3 (-C -R -g -T -u -t -f --ALIGNERS gmap). Predicted genes based on the transcripts were compared to SWISS-PROT Database by BLASTP v2.9 with the minimum identity requirement of 95% and then the top 3,000 hits were selected as the training-sets to train AUGUSTUS v3.3.1 with default parameters. *De novo* prediction evidence, protein homology evidence and transcript evidence were combined and the gene models of the *W. australiana* genome were predicted by EVidenceModeler with parameters "--segmentSize 1000000 --overlapSize 100000". The obtained EVidenceModeler genes were aligned against the TransposonPSI (http://transposonpsi.sourceforge.net/) database to exclude predicted genes containing TE-related domains. Genes with length that could not be divided by 3 were also excluded based on the principle of codon translation. The protein sequences were subsequently mapped to various functional annotation databases by BLASTP v2.9 with parameters “-evalue 1e-5, -max_target_seqs 1”, including Non-Redundant protein sequence databases (NR), Kyoto Encyclopedia of Gene and Genomes (KEGG) database, SwissProt database and Eukaryotic orthgene Groups of protein (KOG) database. The Gene Ontology (GO) analysis was performed by Interproscan v5.32-71.0 (44) with default parameters. To further assess the quality and completeness of the predicted gene model, protein sequences were aligned to the ‘embryophyta_odb10’ dataset of BUSCO v3.1.0 to evaluate the completeness of conserved gene set.

**Genome Phylogenetic Analysis.** *W. australiana* and fourteen other species (Dataset, Table S12) were included for phylogenic evolutionary analysis. The OrthoMCL pipeline v2.0.9 was used to identify gene families between genomes of these species and the protein sequences of the longest transcript of each gene among these species were mutually aligned using BLASTP with parameters "-evalue 1e-5" to obtain the sequence similarity information. Orthologous genes and paralogous genes were subsequently classified. Orthologous single-copy genes were used as the seed sequences to perform the phylogenetic analysis. The single-copy gene sequences were aligned by MAFFT v7.313 and then the above result was trimmed by Gblocks v0.91b with parameters "-t = p -b5 = h" to construct a phylogenetic tree based on the GTRGAMMA model and 1,000 bootstrap replicates by RAxML v8.2.10 (-m PROTGAMMAAUTO -p 12345 -T 8 -f b). Divergence time obtained by TimeTree was set as calibration points (Dataset, Table S13) to divergence time estimation among fifteen species with RelTime by default parameters. Gene family contraction and expansion analysis was performed by CAFE v4.2.1 with parameters "-p 0.05 -t 10 -r 10000", which applies a birth and death rate to model gene family size over a phylogeny based on orthologous groups from OrthoMCL and the ultrametric tree obtained from RelTime. Expanded or contracted gene families with viterbi p <= 0.05 and family-wise p <= 0.01 were defined as significant compared with the last common ancestor. Both GO and KEGG enrichment analyses of significant gene families were performed by clusterProfiler Package (45) with parameters “pAdjustMethod = 'BH', pvalueCutoff = 1, qvalueCutoff = 1”.

**RNA Extraction and Library Preparation for Single-plant RNA Sequencing.** Single-plant *Wolffia* RNA was extracted using the protocol of RNeasy® plant mini kit (Qiagen cat# 74903). RNA concentration was measured using a Qubit RNA assay kit (ThermoFisher, Ref# Q32855A) in a Qubit 2.0 fluorometer (Life Technologies, Ref# REQ32866). Total RNA of 100 ng per sample was used as input material for RNA sample preparations.

Libraries were generated using a NEBNext Ultra$^{TM}$ RNA library prep kit Illumina (New England Biolabs, Ipswich, MA, USA, Cat# E7530) and index codes were added to attribute sequences to each sample. The mRNA was purified from total RNA using poly-T oligo-attached magnetic beads (New England Biolabs, Ipswich, Massachusetts, USA, Cat# E7490). First-strand cDNAs were synthesized by the NEBNext RNA First Strand Synthesis Module

(New England Biolabs, Ipswich, MA, USA, Cat# E7525L). Second-strand synthesis was performed by the NEBNext Ultra II Non-Directional RNA Second Strand Synthesis Module (New England Biolabs, Ipswich, MA, USA, Cat# E6111L). Library was prepared using KAPA Hyper Prep Kit (KAPA Biosystems, Wilmington, Massachusetts, USA, Cat# KK8504). Products were purified with AMPure XP system (Beckman Coulter, Cat# A63882), and library quality was assessed using the Agilent Bioanalyzer 2100 system (Agilent Technologies, Ref# G2939BA). The library was sequenced on the Illumina HiSeq 4000 platform.

**Analysis of Genes Related to Morphological Processes.** The *Arabidopsis thaliana* root and flower genes (Dataset, Table S14) were used as the seed sequences to align against the *W. australiana*'s gene set. Meanwhile, we used the same method to the gene set of *Oryza sativa* (Dataset, Table S16). Then, these genes were classified to different gene families based on the previous research of *A. thaliana.* Next, each of gene family was used to construct gene tree by MEGAX (sequences aligned by MUSCLE and tree constructed by maximum likelihood method).

**Differential Expression Genes (DEGs) Analysis.** Different datasets of transcriptome (Control, C; Induced, I; and Flowered, F) of the *W. australiana* were obtained (Dataset, Table S1). Based on the different transcriptome datasets, read counts of genes in each sample obtained from hisat2 and StringTie with default parameters, DEGs among different sample comparisons were identified by DESeq2 Package (46). Genes with a false discovery rate (FDR) $\leq 0.05$ and fold-change $\geq 2$ were considered to be DEGs in each comparison (Dataset, Table S17). Both GO and KEGG enrichment analysis of DEGs in each group was performed using clusterProfiler Package with parameters "pAdjustMethod = 'BH', pvalueCutoff = 1, qvalueCutoff = 1".

**Chromosome FISH Analysis.** Chromosome preparation was conducted according to a previous study (47). The entire plants were harvested and pretreated in 0.002 M 8-hydroxyquinoline at 23 °C for 2 h, and then fixed with Cannoy Solution (methanol: acetic acid = 3:1) at 25 °C overnight. After washed with distilled water, the plants were digested using the enzyme mixture of 2% cellulase, 1% pectinase at 37 °C for 90 min. After that, they were quickly rinsed in distilled water and fixed with the same fixation solution. The samples were spread on precooled slides and dried by passing through the flame of a Bunsen burner 3 or 4 times. Chromosomes were counterstained with 4',6-diamidino-2-phenylindole (DAPI) in an anti-fade solution (Vector Laboratories, Burlingame, CA, USA). The FISH assay was conducted using 45S rDNA as a probe labeled with digoxigenin and detected by anti-digoxigenin-rhodamine (48). Images were captured under a Zeiss A2 fluorescence microscope with a micro-charge-coupled camera (Zeiss, Germany).

**Microscopy.** The *W. australiana* and *S. polyrhiza* plantlets were macerated with 10% peracetic acid at 80 °C for 30 min. After extensive washing, the plantlets were squashed and stained with cell wall dye Direct Red 23 (0.01% v/v) and photographed using a fluorescence microscope (Imager D2, Zeiss).

To test the function of the only SCW-related VND homolog *WausLG14.977* in vessel differentiation, the full-length coding sequence of *WausLG14.977* was cloned and inserted into a binary vector pCAMBIA1300 between *CaMV 35S* promoter and *NOS* terminator. The resulting construct was transfected into *A. tumefaciens* strain EHA105 and infiltrated into one-month-old tobacco leaves. Autofluorescence signals of the induced vessel wall were recorded with 405 nm excitation using a laser scanning confocal microscope (LSM 980, Zeiss).

**Cell Wall Composition Analysis.** The *W. australiana* plantlets were collected, freeze-dried, and subjected for cell wall composition determination as described previously (49). In brief, the alcohol insoluble residues were prepared by ball milling and successive extraction, and then incubated with α-amylase (Sigma) at 97 °C for 35 min and 60 °C for 1 h to remove starch. The destarched alcohol insoluble residues were mildly hydrolyzed using 2 M trifluoroacetic acid. The supernatants were treated with sodium borohydride solution to reduce alditols and then with acetic anhydride to acetylate. The resulting alditol acetates were extracted with ethyl acetate and quantified using gas chromatography (7890, Agilent)-coupled mass spectrometry (5977, Agilent). The pellets remained were treated in Updegraff reagent (acetic acid:nitric acid:water, 8:1:2, v/v) at 100 °C for 30 min and further hydrolyzed with 72% (v/v) sulfuric acid. Cellulose content was measured via anthrone assays. Five biological replicates were included in the examinations.

**Phylogenetic Analysis.** A neighbor-joining phylogenetic tree of SCW-related NAC homologs in *W. australiana*, *P. patens*, *S. moellendorffii*, *A. trichopoda*, *O. sativa* and *A. thaliana* was built using MEGA6 software (50) with 1000 bootstrap replicates. The sequences were obtained from the online server PLAZA (https://bioinformatics.psb.ugent.be/plaza/), except that the sequences of *W. australiana* homologs were obtained from this study. Sequence alignment of WausLG14.977 with the NAC homologs, AtSND1, OsSNAC1 (3ULX), AtANAC (1UT7) and AtANAC019 (3SWM), was conducted using ClustalW (https://www.clustal.org) and ENDscript/ESPript (https://endscript.ibcp.fr).

The 3D protein structures of WausLG14.977 and AtSND1 (At1g32770) were predicted by using the RoseTTAFold server (https://robetta.bakerlab.org) according to the instructions (51). The predicted Model 1 was displayed and applied for structure comparison using UCSF Chimera (https://www.rbvi.ucsf.edu/chimera). The protein structures of NAC homologs SNAC1 (3ULX), ANAC (1UT7) and ANAC019 (3SWM) were downloaded from RCSB PDB database (https://www.rcsb.org/). Z-scores of WausLG14.977 protein structures to that of Arabidopsis SND1 and rice SNAC1 were determined using the Dali server (https://ekhidna2.biocenter.helsinki.fi/dali).

**Transcription Regulatory Network (TRN) Analysis.** To build the regulatory network, we first called putative Transcription Factors (TFs) as well as corresponding binding motifs in *W. australiana* genome based on the established pipelines (25), then scanned the promoter regions of all predicted *W. australiana* genes (TSS −500 bp to +100bp) with the identified motifs systematically (by FIMO, cut-off 1e-5) to connect the TFs and their potential targets. The inferred network was further refined via transcriptomic data with the following criteria: a) the TF-target pair should be co-expressed (Spearman correlation > 0.9); b) at least one gene in the TF-target pair should be differentially expressed between control samples and flowered samples or induced but not flowered samples, c) at least one gene in the TF-target pair should be the floral organ initiation or differentiation related genes (Dataset, Table S19).

**Protein Structure Prediction and Classification.** The predicted mRNAs were firstly translated into protein sequences. MMseqs2 (32) program suite was employed for further protein sequence analysis, cluster submodule was then used for clustering to reduce redundancy of sequence space, with e-value threshold set to 1e-5. Representative sequences of clusters were compared with those available in structure database (*i.e.*, PDB (52, 53) and AlphaFold Protein Structure Database (31) using search submodule, with e-value threshold set to 1e-5 as well). As a result, 6800 non-homologous protein sequences were left for further structural prediction.

Non-docker version AlphaFold2 (30) was deployed for speed and scalability. Features (*i.e.*, multiple sequence alignments) needed as input for further prediction were firstly generated on a distributed cluster of machines without GPUs. Further structural prediction by neural network and refinement using molecular dynamics were both conducted on machines with graphics cards. Each task was provided with one graphics card to speed up computation. Finally, we obtained 6,798 predicted structures and their relative information, while the prediction for the other two failed due to video memory limitation.

To identify the superfamily of these predicted structures, we used DaliLite.v5 (54) (*i.e.*, a standalone program for protein structural alignment using Dali method) to compare these with representative structures of superfamilies provided by SCOPE (55, 56). The all-against-all structural comparisons were performed with default parameters. The hits with the highest Z score were considered as the best ones, and thus the superfamilies of query predicted proteins were considered as same as those of best hits.

**Single Nucleus Isolation, Single-Nucleus Library Construction and Sequencing, Raw Data Processing, and Generation of Gene Expression Matrix.** Fresh samples of *W. australiana* were used for single nucleus isolation as described previously (27, 57) with slight modifications. The single-nucleus library construction and sequencing were conducted as previously described (58). The DNBelab C Series Single-Cell Library Prep Set (MGI, Cat# 1000021082) was utilized as in (59).

Raw reads were demultiplexed and mapped to the Waus reference genome by PISA (Version 1.1.0). The pipelines include align reads and generate gene-cell matrices. STAR was used with default parameters for mapping and transcript quantifications. The gene-cell matrices generated by R package Seurat were used for subsequent analysis.

**Cell Clustering, Cell Type Identification, Quality Control and Cell Type Annotation for snRNA-seq.** Downstream analyses were mainly performed with Seurat. The gene-cell matrices of 42,667 cells were loaded into the Seurat package implemented in R (Version 3.6.3). The analysis was performed as previously described (58).

To filter out the low-quality cells, the following quality control criteria were followed: one gene is expressed in ⩾ three cells, and one single cell expresses ⩾ 200 genes. We filtered the cells with gene numbers > 3000 or < 200, and unique gene counts < 500. After applying these quality control criteria, 15,983 cells and 14,812 genes were kept for subsequent analysis.

The cell type of each cell cluster was manually defined by the high-expressed genes in each cell cluster, based on these cluster-enriched genes' correspondence to Arabidopsis genes. The Arabidopsis genes' expression information was retrieved from BAR eFP Browser (60) from TAIR (https://www.arabidopsis.org/index.jsp).

**Gene Transformation of *W. auatraliana*.** The *Agrobacterium tumefaciens* strain LBA4404 harboring the plasmids of synthetic reporters -TCS::GUS/pKGWFS7.0 which have been used to report cytokinin output was used in our study (34). Approximately 0.5 g fresh *Wolffia* and 1 g sterilized glass beads (1 mm) were placed in 2-ml sterilized Eppendorf microcentrifuge tube with 500 µL *Agrobacterium* suspension, 1 µL silwet L-77 and 100 µM Acetosyringone (AS). The tubes were first subjected to ultrasound at a frequency of 40 kHz for 1 min at 28 °C. After sonication treatment, a vacuum of approximately 0.7 $kg/cm^2$ was applied for 10 min. Then the tubes were shaken around 150 rpm for 15 min at 28 °C. Finally, plants were transferred onto filter papers wetted with liquid half SH medium adding 1% sucrose (w/v) and 100 µM AS at pH 5.2 in the dark for 5 d at 25 °C.

After five days of co-cultivation, the infected *Wolffia* plants were transferred to selection medium: solid half SH medium supplemented with 300 mg/L cefotaxime, 20 mg/L G418 (Geneticin), and 1% (w/v) sucrose under a 16 h/8 h photoperiod of approximately 85 µmol $m^{-2}$ $s^{-1}$ of white light at least for 4 weeks. Then the survived plants were transferred to induction medium: solid half SH medium adding 150 mg/L cefotaxime, 20 mg/L G418 (Geneticin) and 1% (w/v) sucrose. The resistant plants were identified by the β-Glucuronidase (GUS) assay. The transformation efficiency was calculated as percentage of GUS positive plants in the total number of explants. The highest transformation efficiency in current study was approximately

46%.

Note: all materials are listed in *SI Appendix*, Supplementary Table S-Materials (61–82).

**Statistical Analysis.** All details of the statistics applied are provided alongside in the materials and methods, figures and tables, and the corresponding legends.

**Supplementary Figures**

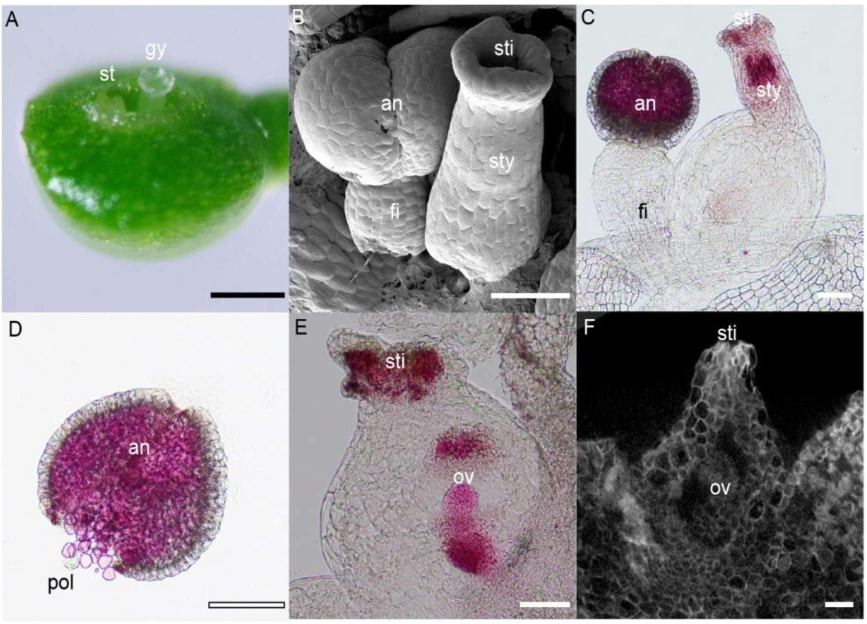


**Fig. S1.** The *Wolffia australiana* floral organ structure. The *Wolffia australiana* floral organ is shown in (*A*)-(*F*). Abbreviations: st, stamen; gy, gynoecium; an, anther; fi, filament; sti, stigma; sty, style; pol, pollen; ov, ovule. Bars: black = 1 mm, white = 100 μm.

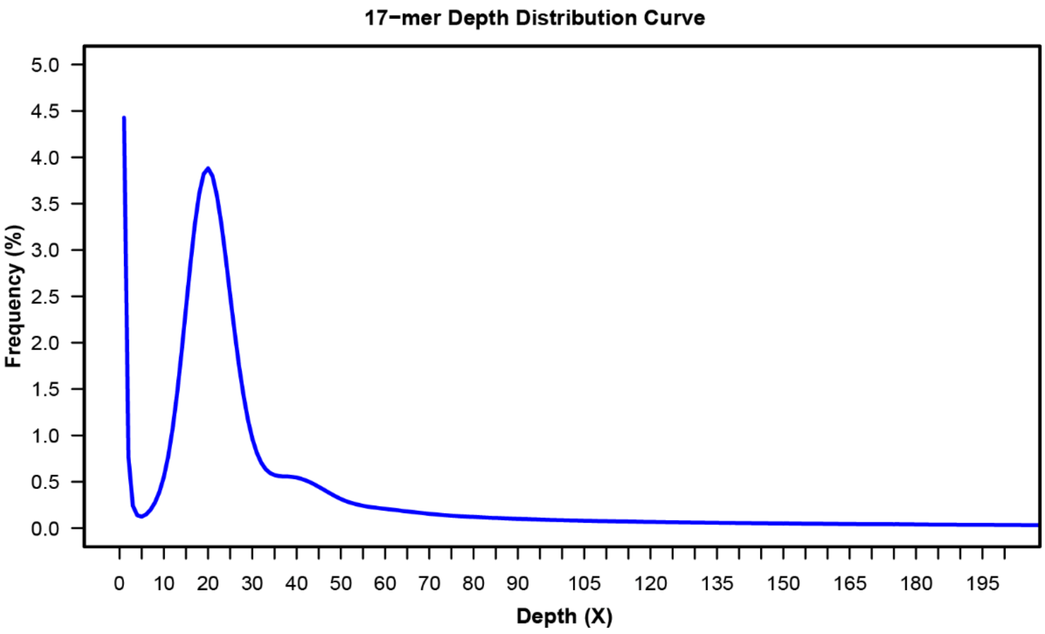


**Fig. S2.** The frequency distribution of 17-mer.

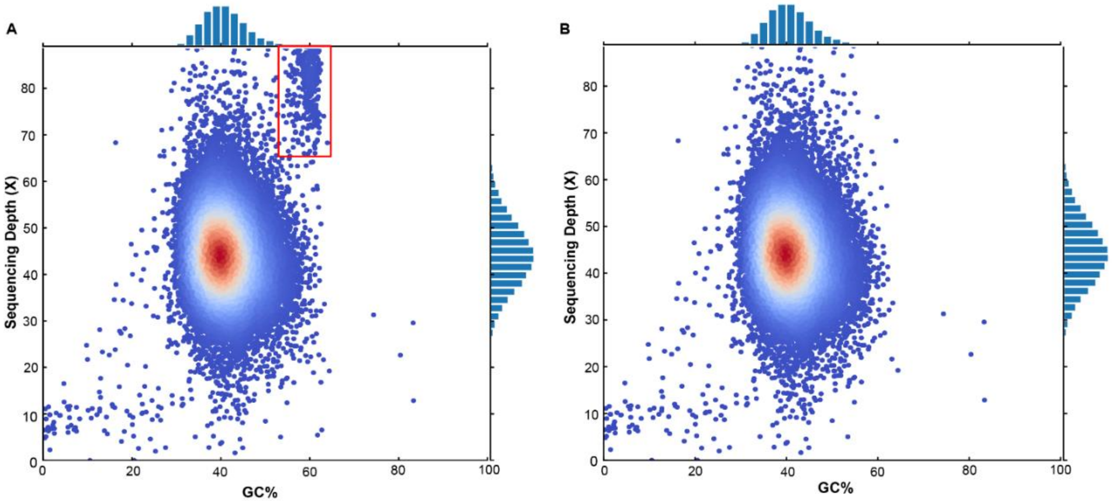


**Fig. S3.** The GC-Depth of *W. australiana* genome based on the 10-kb bins. Panel (*A*) is the genome version G2, and panel (*B*) is the genome version G3. These bins of contaminative contigs are in the red frame.

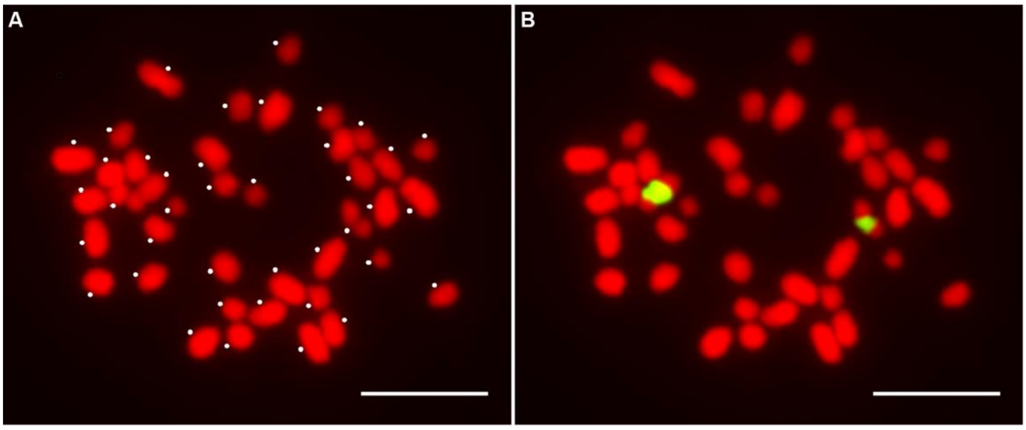


**Fig. S4.** Somatic metaphase chromosomes of *W. Australiana*. Chromosomes (pseudocolored in red) were stained with 4′,6-diamidino-2-phenylindole (DAPI). (*A*) The white dot beside each chromosome indicates the individual chromosome, and the total chromosome number was 40. (*B*) The same cell probed with 45S rDNA showing a pair of chromosomes with 45S rDNA (pseudocolored in green) located in the middle of the chromosome. Bar = 5 μm.

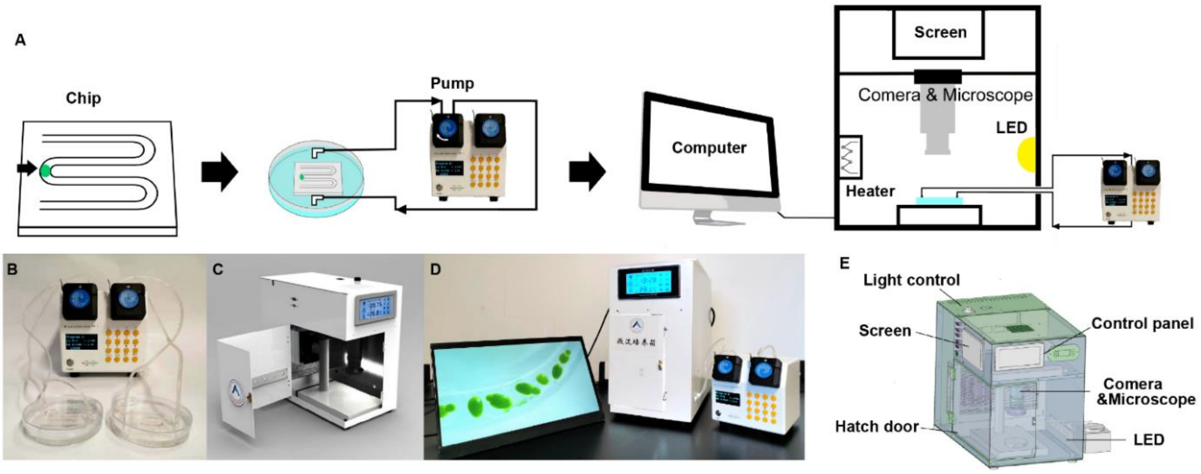


**Fig. S5.** Composition of the Plant-on-Chip Culture System. (*A*) Schematic structure of the system. (*B*) The chip and pump. (*C*) Design sketch of the incubator. (*D*) The PoC system. (*E*) The blueprint of the incubator.

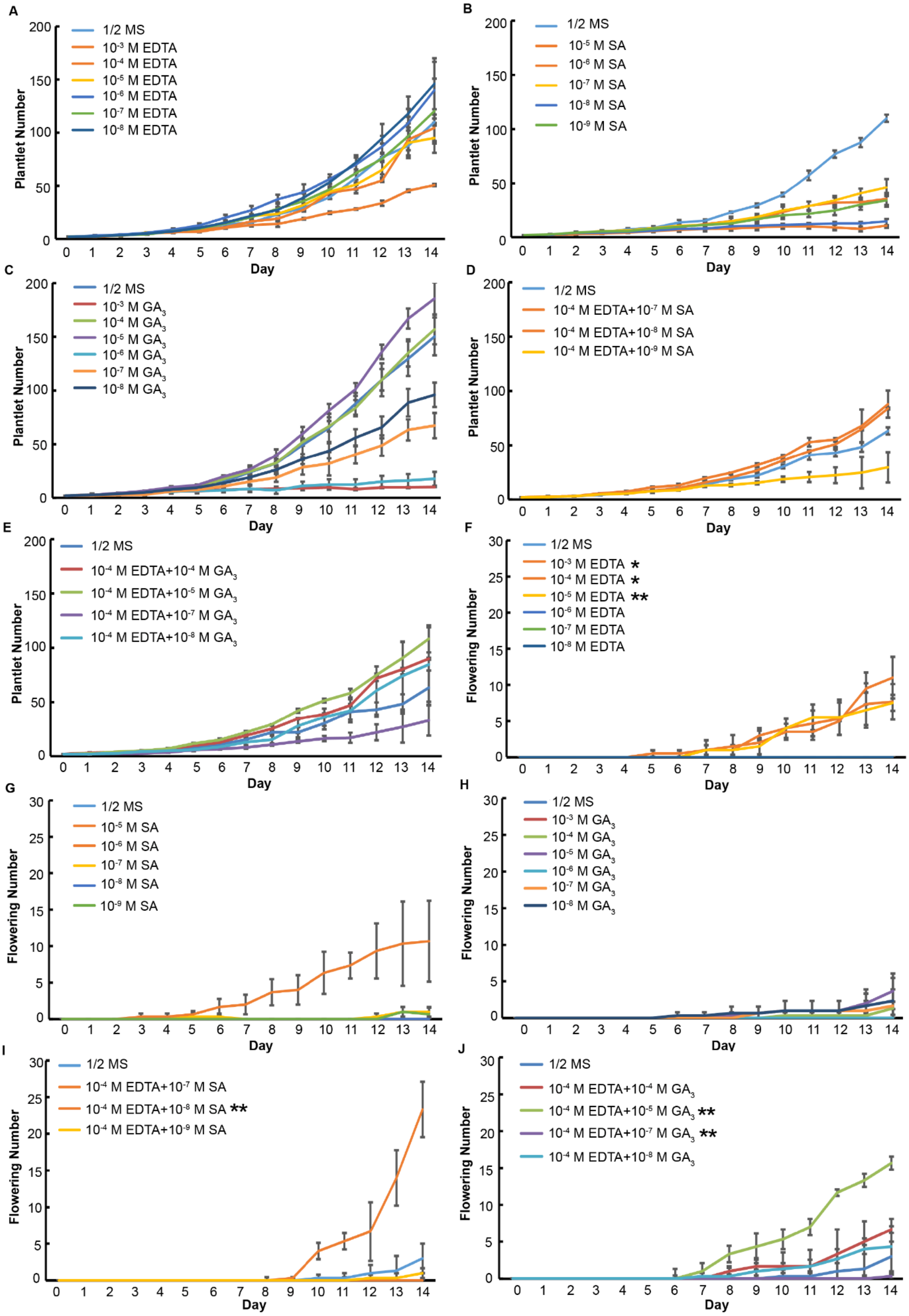


**Fig. S6.** The *W. australiana* Flowering Rate upon Various Treatments. The *W. australiana* branching speed is shown in (*A*)-(*E*)€. The plantlet was grown on 1/2 MS medium with EDTA

(*A*), SA (*B*), $GA_3$ (*C*), EDTA+SA (*D*), or EDTA+$GA_3$ (*E*) for 2 weeks. The flowering plantlet numbers of each group are shown in *F-J* (mean ± SEM, n = 3) *$P < 0.05$, **$P < 0.01$, two-tailed Student's *t*-test for all pairwise comparisons.

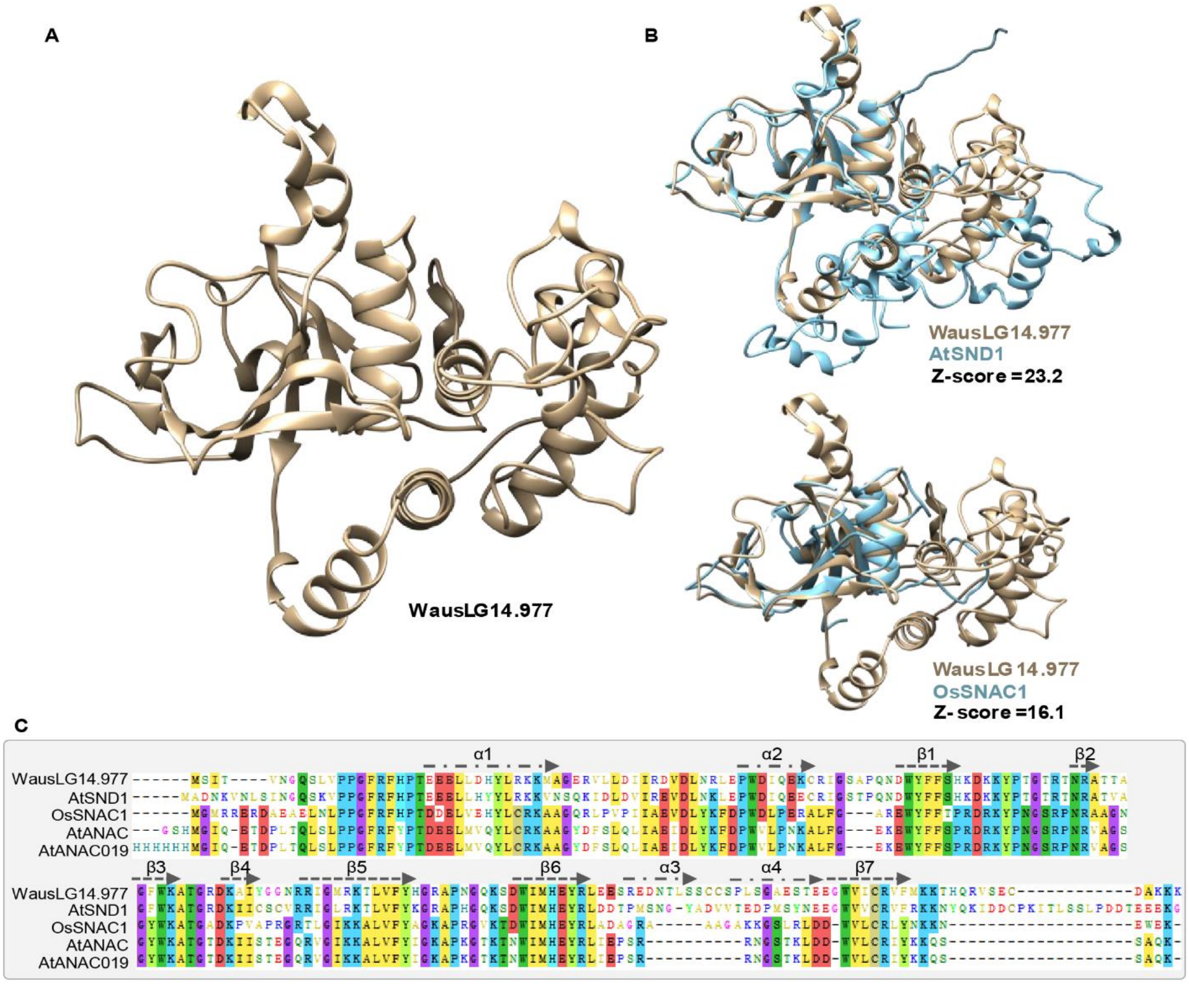


**Fig. S7.** Protein structure analyses of WausLG14.977 predicted using RoseTTAFold. (*A*) The 3D structure of WausLG14.977 predicted by using the RoseTTAFold server (https://robetta.bakerlab.org). (*B*) Structure comparison of WausLG14.977 with that of AtSND1 and OsSNAC1. Z-scores are indicated. (*C*) Sequence alignment of WausLG14.977 and the NAC homologs using ClustalW and ENDscript/ESPript. Z-scores of WausLG14.977 protein structure relative to those of Arabidopsis SND1 and rice SNAC1 were determined using the Dali server (https://ekhidna2.biocenter.helsinki.fi/dali).

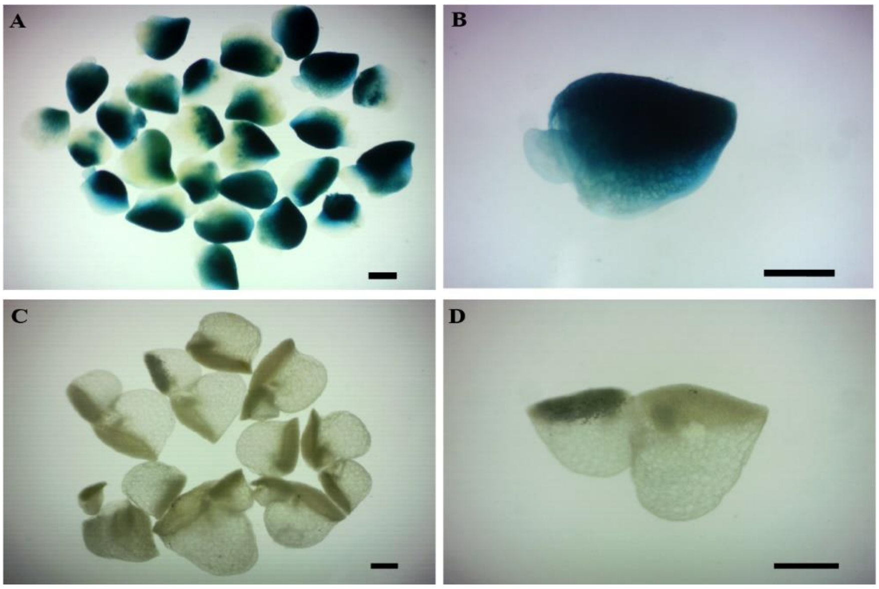


**Fig. S8.** Transgenic plantlets with GUS staining of *W. australiana*. (*A*-*B*) GUS staining showing the expression of cytokinin in *W. australiana.* (*C*-*D*) GUS staining of Wolffia transformed by an empty vector. Bar = 500 µm.

**Table S-Materials. All reagents, data and software are listed below**

| REAGENT or RESOURCE | SOURCE | IDENTIFIER |
|---|---|---|
| **Chemicals, Peptides, and Recombinant Proteins** | | |
| Acetosyringone | Sigma-Aldrich | CAS 2478-38-8 |
| Cefotaxime | Sigma-Aldrich | CAS 64485-93-4 |
| Geneticin | Sigma-Aldrich | CAS 108321-42-2 |
| X-Gluc | Sigma-Aldrich | CAS 129541-41-9 |
| Silwet-77 | Coolaber | Cat# CS9791 |
| Peracetic acid | Aladdin | Cat# P299577 |
| Direct Red 23 | Sigma-Aldrich | Cat# 212490 |
| Trifluoroacetic acid | Sigma-Aldrich | Cat# T6508 |
| α-Amylase | Megazyme | Cat# E-BLAAM |
| Sodium borohydride | Sigma-Aldrich | Cat# 213462 |
| Sucrose | Sigma-Aldrich | Cat# S8501 |
| Glycerol | Sigma-Aldrich | Cat# G5516 |
| Ethyl acetate | J.T.Baker | Cat# 9282-03 |
| Anthrone | Sigma-Aldrich | Cat# 319899 |
| Acetic acid | J.T.Baker | Cat# 9515-03 |
| Glutaraldehyde | Sigma-Aldrich | Cat# G7651 |
| Paraformaldehyde | Sigma-Aldrich | Cat# P6148 |
| Formaldehyde | Sigma-Aldrich | Cat# 252549 |
| Glycine | Sigma-Aldrich | Cat# 241261 |
| DpnII | New England Biolabs | Cat# R0543S |
| Biotin-14-dCTP | Thermo Fisher Scientific | Cat# 19518018 |
| T4 DNA polymerase | New England Biolabs | Cat# M0203S |
| Cellulase | Sigma-Aldrich | Cat# C0615 |
| Pectinase | Sigma-Aldrich | Cat# P2611 |
| Trichloroacetaldehyde | LMAI Bio | Cat# LM008072 |
| TRIzol reagent | Invitrogen | Cat# 15596018 |
| CTAB | Sigma-Aldrich | Cat# H6269-100G |
| LiCl | Sigma-Aldrich | Cat# L9650 |
| Thiocarbohydrazide | Sigma-Aldrich | Cat# 223220 |
| Digoxigenin | Sigma-Aldrich | Cat# D9026 |
| Anti-digoxigenin-Rhodamine | Sigma-Aldrich | Cat# 11207750910 |
| Sodium borohydride | Sigma-Aldrich | Cat# 213462 |
| Acetic anhydride | Sinopharm | Cat# 10000318 |
| Anthrone | Sigma-Aldrich | Cat# 319899 |
| Ethyl acetate | Sigma-Aldrich | Cat# 270989 |
| $Na_2HPO_4 \cdot 12H_2O$ | Sigma-Aldrich | Cat# 71649 |
| $NaH_2PO_4 \cdot H_2O$ | Sigma-Aldrich | Cat# 71507 |
| Osmium Tetroxide, 4% aqueous ($OsO_4$) | Ted Pella | Cat# 18459 |
| Potassium ferrocyanide | Sigma-Aldrich | Cat# P3289 |

| | | |
|---|---|---|
| 8-hydroxyquinoline | Sigma-Aldrich | Cat# 252565 |
| Methanol | Sigma-Aldrich | Cat# 1424109 |
| Nitric acid | Sinopharm | Cat# 10014518 |
| Sulfuric acid | Sigma-Aldrich | Cat# 339741 |
| Anti-fade solution | Vector Laboratories | Cat# H-1200-10 |
| Spurr's resin | SPI Supplies | Cat# 02680-AB |
| Poly-T oligo-attached magnetic beads | New England Biolabs | Cat# E7490 |
| MS medium | Solarbio | Cat# M8521 |
| SH medium | Solarbio | Cat# LA8620 |
| $GA_3$ | Aladdin | Cat# G105689 |
| SA | Sigma | Cat# S7401 |
| EDTA | Sigma | Cat# E4884 |
| PDSM | Momentive | Cat# RTV615 |
| Pump | MesoBioSystem | Cat# pp1 |
| Critical Commercial Assays | | |
| RNeasy® Plant Mini Kit | Qiagen | Cat# 74903 |
| Qubit RNA Assay Kit | Life Technologies | Cat# Q32852 |
| NEBNext Ultra™ RNA Library Prep Kit Illumina | New England Biolabs | Cat# E7530L |
| NEBNext First Strand Synthesis Module | New England Biolabs | Cat# E7525L |
| NEBNext Ultra II Non-Directional RNA Second Strand Synthesis Module | New England Biolabs | Cat# E6111L |
| Bionano Prep DLS Labeling DNA Kit | Bionano Genomics | Cat# 80005 |
| Bionano Prep DLS Labeling DNA Kit | Bionano Genomics | Cat# 80005 |
| KAPA Hyper Prep Kit | KAPA Biosystems | Cat# KK8504 |
| TruSeq RNA Library Preparation Kit | Illumina | Cat# RS-122-2001/2002 |
| M-MuLV reverse transcriptase | New England Biolabs | Cat# M0253L |
| USER enzyme | New England Biolabs | Cat# M5505L |
| AMPure XP system | Beckman Coulter | Cat# A63882 |
| Single-Cell Library Prep Set | MGI | Cat# 1000021082 |
| Ligation Sequencing Kit | Nanopore store | Cat# SQK-LSK109 |
| **Deposited Data** | | |
| *Wolffia australiana 7733* (Waus) | *Wolffia australiana 7733* (Waus) genome, this study | NCBI Genome: CP092600-CP092619, 20 chromosomes. NCBI sequence read archive (SRA): PRJNA808652, Nanopore; PRJNA808655, Illumina |

| | | |
|---|---|---|
| | | genome; PRJNA808685, Hi-C; PRJNA808734, BioNano; PRJNA808736, RNA-seq for genome; PRJNA808739, single-plant RNA-seq; PRJNA809022 single-nucleus RNA-seq. NCBI Supplementary Files: SUPPF_0000004267, BioNano. http://wolffiapond.net/ |
| *Chlamydomonas reinhardtii* | *Chlamydomonas reinhardtii* genome | https://www.ncbi.nlm.nih.gov/genome/?term=Chlamydomonas+reinhardtii+ |
| *Azolla filiculoides Lam.* | *Azolla filiculoides Lam.* genome | https://www.fernbase.org/?tdsourcetag=s_pcqq_aiomsg |
| *Physcomitrella patens* | *Physcomitrella patens* genome | https://www.ncbi.nlm.nih.gov/genome/?term=Physcomitrella+patens |
| *Amborella trichopoda* | *Amborella trichopoda* genome | https://www.ncbi.nlm.nih.gov/genome/?term=Amborella+trichopoda |
| *Oryza sativa L.* | *Oryza sativa L.* genome | https://www.ncbi.nlm.nih.gov/genome/?term=Oryza+sativa |
| *Spirodela polyrhiza* | *Spirodela polyrhiza* genome | https://www.ncbi.nlm.nih.gov/genome/?term=Spirodela+polyrhiza |
| *Spirodela polyrhiza 9509* | *Spirodela polyrhiza 9509* genome | ftp://ftp.lemna.org/spirodela_polyrhiza_9509/ |
| *Lemna minor 8627* | *Lemna minor 8627* genome | ftp://ftp.lemna.org/lemna_minor_8627/ |
| *Lemna gibba 7742a* | *Lemna gibba 7742a* genome | ftp://ftp.lemna.org/lemna_gibba_7742a/ |
| *Wolffia australiana 8730* | *Wolffia australiana 8730* genome | ftp://ftp.lemna.org/wolffia_australiana_8730/ |
| *Colocasia esculenta* | *Colocasia esculenta* genome | https://www.ncbi.nlm.nih.gov/genome/?term=Colocasia+esculenta |
| *Zostera marina* | *Zostera marina* genome | https://www.ncbi.nlm.nih.gov/genome/?term=Zostera+marina |
| *Elaeis guineensis* | *Elaeis guineensis* genome | https://www.ncbi.nlm.nih.gov/genome/?term=Elaeis+guineensis |
| *Arabidopsis thaliana* | *Arabidopsis thaliana* genome | https://www.ncbi.nlm.nih.gov/genome/?term=Arabidopsis+thaliana) |
| *Nymphaea colorata* | *Nymphaea colorata* genome | https://www.ncbi.nlm.nih.gov/genome/69117 |
| *Utricularia gibba* | *Utricularia gibba* genome | https://www.ncbi.nlm.nih.gov/genome/?term=Utricularia+gibba |
| *Cuscuta australis* | *Cuscuta australis* genome | https://www.ncbi.nlm.nih.gov/genome/?term=Cuscuta+australis |
| *Musa acuminata* | *Musa acuminata* genome | http://plants.ensembl.org/Musa_acuminata/Info/Index |
| Kyoto Encyclopedia of Genes and Genomes | Biological organisam pathways database | https://www.kegg.jp/ |

| | | |
|---|---|---|
| Eukaryotic orthgene Groups of protein | Eukaryotic coding proteins' phylogenetic classification database | http://genome.jgi-psf.org/help/kogbrowser.jsf |
| SwissProt | Validated rigorously de-redundant protein sequences database | https://web.expasy.org/docs/swiss-prot_guideline.html |
| Non +72:75redundant Protein sequence databases | Non redundant Protein sequence database | ftp://ftp.ncbi.nlm.nih.gov/blast/db/ |
| PDB sequence data | (61, 62) | ftp://ftp.wwpdb.org/pub/pdb/derived_data/pdb_seqres.txt |
| SCOPE 2.08-stable | (63, 64) | https://scop.berkeley.edu/downloads/ver=2.08 |
| AlphaFold Protein Structure Database: proteome-wide predictions | (65) | https://www.alphafold.ebi.ac.uk/download#proteomes-section |
| **Experimental Models: Organisms/Strains** | | |
| *Wolffia ausriliana* | Institute of Hydrobiology, CAS | Accession: wa7733 |
| **Oligonucleotides** | | |
| Primer: LG14-GW-F. GGGGACAAGTTTGTACAAAAAAGCAGGCTACATGAGCATCACGGTCAACG | This study | N/A |
| Primer: LG14-1-R. TGACCCAATCCTGCACTTCTCTTGAATGTCCCAAGGCTCAA | This study | N/A |
| Primer: LG14-2-F.AGAAGTGCAGGATTGGGTCA | This study | N/A |
| Primer: LG14-2-R.CCGGAAAGCGGACTACAACAACTGGAAAGGGTGTTATCTTCT | This study | N/A |
| Primer: LG14-3-F.TGTTGTAGTCCGCTTTCCGG | This study | N/A |
| Primer: LG14-GW-R.GGGGACCACTTTGTACAAGAAAGCTGGGTACTATCCACCGGGTGTAGA | This study | N/A |
| Recombinant DNA | | |
| pCAMBIA1300-PacI | (66) | Addgene ID: 44183 |
| pCAMBIA1300-WausLG14.977 | This study | N/A |
| **Software and Algorithms** | | |

| | | |
|---|---|---|
| AlphaFold v2.0.1 | (67) | https://github.com/deepmind/alphafold |
| MMseqs2 Release 13-45111 | (68) | https://github.com/soedinglab/MMseqs2 |
| DaliLite.v5 | (69) | http://ekhidna2.biocenter.helsinki.fi/dali/ |
| Pandas 1.3.3 | (70) | https://zenodo.org/record/5501881#.YckwaGBBxhE |
| STAR V2.7.4a | (71) | |
| PISA V0.8.2 | (72) | https://github.com/shiquan/pisa |
| Seurat V3.2.1 | (73) | https://satijalab.org/seurat |
| Seurat V3.2.2 | (73) | https://satijalab.org/seurat |
| Seurat V3.6.3 | (73) | https://satijalab.org/seurat |
| MEGA6 | (74) | https://www.megasoftware.net/ |
| PLAZA | (75) | https://bioinformatics.psb.ugent.be/plaza/ |
| ClustalW | (76) | https://www.clustal.org/ |
| ENDscript/ESPript | (77) | https://endscript.ibcp.fr |
| RoseTTAFold server | (78) | https://robetta.bakerlab.org/ |
| UCSF Chimera | (79) | https://www.rbvi.ucsf.edu/chimera |
| RCSB PDB database | (62) | https://www.rcsb.org/ |
| Dali server | (69) | https://ekhidna2.biocenter.helsinki.fi/dali |
| BWA 0.7.12-r1039 | (80) | https://sourceforge.net/projects/bio-bwa/ |
| Hmmer 3.0 | (81) | http://www.hmmer.org/download.html |
| FIMO 4.11.4 | (82) | https://meme-suite.org/meme/doc/fimo.html |
| Igraph 1.2.7 | R package | https://igraph.org/r/ |
| BLAST v2.9 | Homolougs alignments for nucleotide sequences | https://blast.ncbi.nlm.nih.gov/Blast.cgi |
| NextDenovo v2.0-beta.1 | Self-error correction software of the Nanopore original data | https://github.com/Nextomics/NextDenovo/ |
| NextGraph v2.0-beta.1 | Genome contig assembly software | https://github.com/Nextomics/NextDenovo/ |
| Minimap2 r41 | Sequence alignment software | https://github.com/lh3/minimap2 |
| Racon v1.4.3 | Error correction software | https://github.com/isovic/racon |
| Fastp v0.19.4 | Quality control software for Illumina raw data | https://github.com/OpenGene/fastp |
| BWA 0.7.12-r1039 | Sequence alignment of genomic data on Illumina platform | http://bio-bwa.sourceforge.net/ |
| NextPolish v1.0.5 | Error correction software of NGS data | https://github.com/Nextomics/NextPolish.git |
| SAMtools v1.4 | Analysis tool for sam and bam files | https://github.com/samtools/samtools |
| BCFtools v1.8.0 | Analysis tool for vcf files | https://github.com/samtools/bcftools |

| | | |
|---|---|---|
| HISAT2 v2.1 | Sequence alignment of transcriptome data on Illumina platform | https://daehwankimlab.github.io/hisat2/ |
| BUSCO v3.1.0 | Genome completeness evaluation software by using OrthoDB database | https://busco.ezlab.org/ |
| CEGMA v2 | Genome completeness evaluation software by using core genes | https://github.com/KorfLab/CEGMA_v2/ |
| Bowtie2 v2.3.2 | Sequence alignment of sequences on Illumina platform | http://bowtie-bio.sourceforge.net/bowtie2/index.shtml |
| Bionano Solve™ data analysis software | Bionano optical genome mapping | https://bionanogenomics.com/products/bionano-data-solutions/ |
| LACHESIS | Hi-C scaffolding software | https://github.com/shendurelab/LACHESIS |
| GMATA v2.2 | Simple sequence repeats identification software | https://sourceforge.net/projects/gmata/ |
| Tandem Repeats Finder v4.07b | Tandem repeat sequences identification software | https://tandem.bu.edu/trf/trf.download.html |
| LTR_finder v1.07 | Long terminal repeat retrotransposons identification software | http://tlife.fudan.edu.cn/tlife/ltr_finder/help/single.html |
| LTR_harverst v1.5.10 | Long terminal repeat retrotransposons identification software | http://genometools.org/tools/gt_ltrharvest.html |
| LTR_retriver v1.8.0 | Long terminal repeat library construction software | https://github.com/oushujun/LTR_retriever.git |
| MITE-Hunter v11-2011 | Miniature inverted transposable elements identification software | http://target.iplantcollaborative.org/mite_hunter.html |
| RepeatModeler v1.0.11 | Genome masking and novel transposable elements identification software | https://github.com/Dfam-consortium/RepeatModeler |
| RepeatMasker v1.331 | Repetitive sequences identification software | http://www.repeatmasker.org/ |
| GeMoMa v1.6.1 | Gene prediction software based on homolog proteins | http://www.jstacs.de/index.php/GeMoMa |
| StringTie v1.3.3d | Gene prediction software based on transcriptome data on Illumina platform | https://ccb.jhu.edu/software/stringtie/ |
| PASA v2.3.3 | Transcriptome assembly software | https://github.com/PASApipeline/PASApipeline/wiki |
| TransDecoder | Protein-coding region prediction software | https://github.com/TransDecoder/TransDecoder/wiki |
| AUGUSTUS v3.3.1 | *De novo* gene prediction software | http://augustus.gobics.de/ |
| EVidenceModeler v1.1.1 | Software combines ab intio gene | https://evidencemodeler.github.io/ |

| | | |
|---|---|---|
| | predictions and protein and transcript alignments into weighted consensus gene structures | |
| Infernal v1.1.2 | rRNA, snRNA and miRNA prediction software | http://eddylab.org/infernal/ |
| tRNAscan-SE v2.0 | tRNAs prediction software | http://lowelab.ucsc.edu/tRNAscan-SE/ |
| RNAmmer v1.2 | rRNAs prediction software | http://www.cbs.dtu.dk/services/RNAmmer/ |
| OrthoMCL v2.0.9 | Homolougs alignments for protein sequences | https://orthomcl.org/orthomcl/ |
| MAFFT v7.313 | Multiple sequence alignments of gene families | https://myhits.sib.swiss/cgi-bin/mafft |
| Gblocks v0.91b | Protein multiple sequence alignment software | http://molevol.local/castresana/Gblocks/ |
| RAxML v8.2.10 | Phylogenetic tree construction software | https://github.com/amkozlov/raxml-ng |
| Figtree v1.4.4 | Phylogenetic tree editing software | http://tree.bio.ed.ac.uk/software/figtree/ |
| CAFE v4.2.1 | Gene family contraction and expansion analysis software | https://github.com/hahnlab/CAFE |
| ClusterProfiler | GO and KEGG enrichment software | https://github.com/YuLab-SMU/clusterProfiler |
| McScanX | Collinear block computing software within or between genomes | http://chibba.pgml.uga.edu/mcscan2/ |
| KaKs_Calculator v2.0 | Synonymous substitution rate calculation software | https://sourceforge.net/projects/kakscalculator2/ |
| DESeq2 package | Differential expression genes calculation software | http://bioconductor.org/packages/release/bioc/html/DESeq2.html |
| Interproscan v5.32-71.0 | Gene Ontology analysis software | https://github.com/ebi-pf-team/interproscan/wiki |
| **Other** | | |
| SEM | Thermo Fisher Scientific | Helios NanoLab G3 UC |
| Cryo-Stage | Quorum Technologies | PP3010T |
| MicroCT | Zeiss | Xradia Context |
| MicroCT | Bruker | SkyScan 1272 |
| Ultramicrotome | Leica Microsystem | UC7 |
| TEM | Jeol | JEM-1400 |
| Qubit 2.0 fluorometer | Life Technologies | REQ32866 |
| Agilent Bioanalyzer 2100 system | Agilent | G2939BA |
| Promethion | Oxford Nanopore Technologies | |
| Illumina NovaSeq 6000 | Illumina | Cat# 20012850 |

| |
|---|
| platform |
